# Simultaneous multicolor DNA-PAINT without sequential fluid exchange using spectral demixing

**DOI:** 10.1101/2021.11.19.469218

**Authors:** Niclas Gimber, Sebastian Strauss, Ralf Jungmann, Jan Schmoranzer

**Affiliations:** Advanced Medical Bioimaging Core Facility, Charité-Universitätsmedizin, Berlin, Germany; Faculty of Physics and Center for Nanoscience, Ludwig Maximilian University, Munich, Germany; Max Planck Institute of Biochemistry, Martinsried, Germany

**Author notes:** Correspondence should be addressed to J.S. and to N.G.

**Keywords:** Super-resolution, single-molecule imaging, fluorescence microscopy, single-molecule localization microscopy, DNA-PAINT, multicolor, spectral demixing

## Abstract

Several variants of multicolor single-molecule localization microscopy (SMLM) have been developed to resolve the spatial relationship of nanoscale structures in biological samples. The oligonucleotide-based SMLM approach ‘DNA-PAINT’ robustly achieves nanometer localization precision and can be used to count binding sites within nanostructures. However, multicolor DNA-PAINT has primarily been realized by ‘Exchange-PAINT’ that requires sequential exchange of the imaging solution and thus leads to extended acquisition times. To alleviate the need for fluid exchange and to speed up the acquisition of current multichannel DNA-PAINT, we here present a novel approach that combines DNA-PAINT with simultaneous multicolor acquisition using spectral demixing (SD). By using newly designed probes and a novel multichannel registration procedure we achieve simultaneous multicolor SD-DNA-PAINT with minimal crosstalk. We demonstrate high localization precision (3 – 6 nm) and multicolor registration of dual and triple-color SD-DNA-PAINT by resolving patterns on DNA origami nanostructures and cellular structures.

## INTRODUCTION

Single-molecule localization microscopy (SMLM) became a widely used method to investigate biological nanostructures with unprecedented spatial resolution in fluorescence-based light microscopy ^1–3^. To investigate the spatial relationships between different protein structures on the nanoscale the precision and applicability of multicolor SMLM is of central importance. However, the development of robust multicolor SMLM approaches was challenged by several factors. These include the limited number of well-performing photoswitchable fluorophores, unwanted color channel crosstalk, chromatic aberrations, multicolor registration errors, long acquisition times and an increasing complexity in the required hardware. Over the last decade, several optimized SMLM approaches that overcome at least some of these limitations have been developed.

Recently, DNA point accumulation for imaging in nanoscale topography (DNA-PAINT) has emerged as a promising SMLM method ^4, 5^. DNA-PAINT uses transient binding of freely diffusing fluorescently labeled oligonucleotides (‘imager strands’, or short: ‘imagers’) to complementary DNA oligonucleotides (‘docking strands’) that are linked to the target structure (i.e. antibody against protein of interest). The transient binding of the imager strands produces the required on/off blinking for SMLM. In contrast to dSTORM ^6^, DNA-PAINT does not require a chemical switching buffer and is largely insensitive to photobleaching as the imager strands are constantly exchanged within the imaging volume. By taking advantage of bright organic fluorophores and the controlled binding kinetics of the imager strands, DNA-PAINT can be used to determine the number of binding sites within a biological sample with high spatial resolution ^7^.

Multicolor DNA-PAINT can be achieved by using multiple docking/imager strand pairs with orthogonal sequences. In the recently developed ‘Exchange-PAINT’ method, multiplexing is realized by sequential fluid exchange of the imaging buffer that contains distinct imager strands for each channel ^4^. This approach reaches very high localization precisions and is insensitive to chromatic errors by using the same dye in each acquisition round. Exchange-PAINT, however, relies on the complete exchange of the imaging strand containing buffer in between the acquisition phases of each channel. Each fluid exchange, including the washing steps, is prone to cause extended sample drift through thermal or pressure gradients, and therefore requires precise drift correction and multichannel registration. The combination of long exposure times that are needed to average out the noise from diffusing imager strands and the sequential color channel acquisition lead to long total acquisition periods. By optimizing both DNA sequences and buffer conditions, the acquisition times were shortened by an order of magnitude ^8^. Most recently, the use of concatenated, periodic DNA sequences offered even faster acquisition speed ^9^. However, all these improved multicolor DNA-PAINT approaches are still limited by the fluid exchange that is required for sequential acquisition.

Multicolor DNA-PAINT without fluid exchange has recently been achieved by modulation of the excitation frequency ^10^. However, this approach (fm-DNA-PAINT) requires a minimum of three rapidly triggered lasers and suffers from chromatic aberrations like in any other multicolor method with spectrally separate fluorophores. Compared to DNA-PAINT, fm-DNA-PAINT was limited to two color channels with a compromised spatial resolution. Another DNA-PAINT variant used three spectrally different dyes and different binding frequencies to achieve sequential multi-color DNA-PAINT without fluid exchange ^11^. This highly multiplexed approach massively speeds up fluorescence in situ hybridization, however, it relies on separable point-like objects and cannot resolve dense cellular structures.

We recently developed a robust multicolor dSTORM approach that achieved high localization precision with fast, simultaneous dual-color acquisition based on ‘spectral demixing’ (SD) ^12–15^. The SD-mode uses spectrally overlapping fluorophores excited by a single laser line and a simple dichroic-based emission splitter to image short and long wavelength components of the emission on two sides of the same camera (Figure 1a). By using a custom open-source software tool (SD-Mixer2 ^16^) the single-molecule localizations are ‘paired’, while non-paired localizations, from which a large part is random noise, are excluded (Figure 1b). Depending on their emission spectra, localizations from each dye display a spatially distinct population within the 2D intensity histogram of long and short channel intensity values (Figure 1c). By applying binary masks to exclude or include populations of the intensity distribution, the colors are assigned to each paired localization before multicolor rendering (Figure 1d). These color separation masks are optimized for maximal inclusion of localizations and minimal crosstalk between the channels.

**Figure 1.**
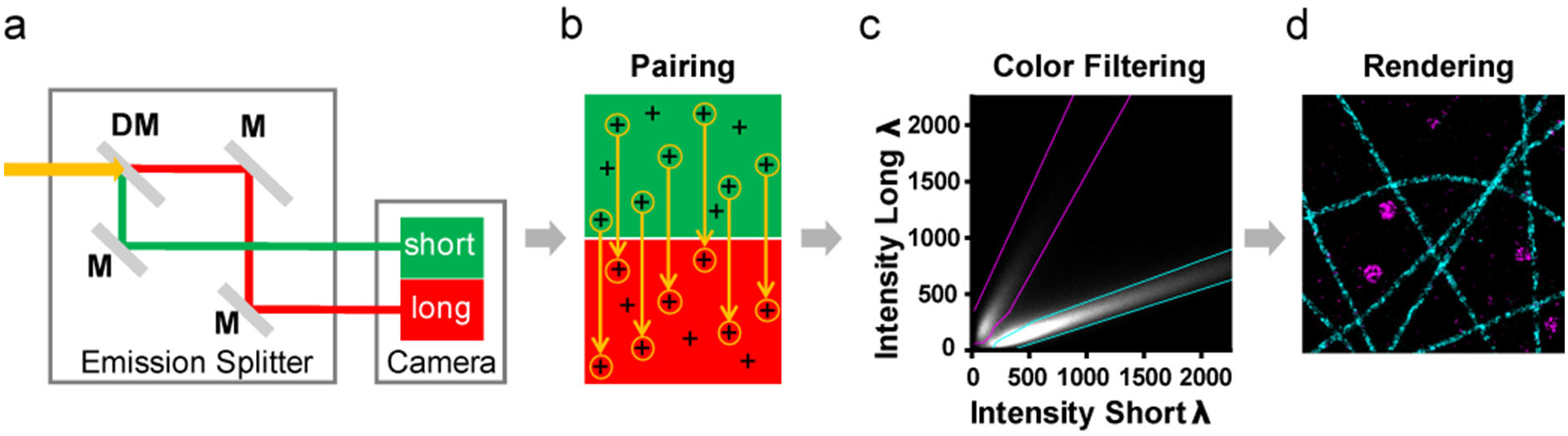
Spectral Demixing (SD) DNA-PAINT principle and workflow. (**a**) Schematic beam path of the emission splitter installed between microscope body (not displayed) and camera system (indicated). All dyes (here dual-color example with ATTO 655 and 700) are excited by a single laser line (e.g. 647 nm) and the mixed emission (yellow) is split via a dichroic mirror (DM) and 100 % mirrors (M) into ‘short’ (green) and ‘long’ (red) wavelength channels onto a single camera. (**b**) The localizations detected in both channels are subjected to a custom ‘pair-finding’ algorithm that identifies corresponding localization pairs. Random (un-paired) localizations are excluded. (**c**) The channel-specific intensity values (short / long) of all localization pairs are plotted into a 2D intensity histogram. Color specific masks (cyan / magenta dotted lines) are designed to minimize color crosstalk and maximize inclusion of localizations. Each included localization is assigned to a color channel according to the masks. (**d**) Example of a rendered dual-color SD-DNA-PAINT image of microtubules (cyan) and clathrin coated vesicles (magenta).

To take advantage of the benefits of DNA-PAINT, but to avoid the need of fluid exchange for multicolor imaging, we here present a novel approach that combines spectral demixing (SD) with DNA-PAINT. In ‘SD-DNA-PAINT’ multiple channels are acquired simultaneously, not sequentially. Compared to Exchange-PAINT, SD-DNA-PAINT effectively speeds up the acquisition time by x-fold for an x-color experiment while keeping the benefits. By avoiding fluid exchange, sample drift and potential errors in multichannel registration are minimized. Using a new combination of spectrally overlapping dyes (ATTO 655, ATTO 680, ATTO 700), we achieved simultaneous triple-color SD-DNA-PAINT with maximal localization precision and minimal crosstalk. We demonstrate the feasibility of SD-DNA-PAINT by resolving known nanostructures within DNA origamis and cells.

## RESULTS AND DISCUSSION

To perform robust multicolor SD-DNA-PAINT we first established a combination of organic dyes suited for this approach. In principle, the choice of dye is not limited to any specific spectral region. The far-red emitting ATTO 655 dye was successfully used to perform DNA-PAINT using a 647 nm laser ^5^. The use of far-red dyes offers the flexibility to add additional non-super-resolved channels with dyes excited in the visible range (405 nm, 488 nm, 561 nm) to the SD-based experiment ^13^. We therefore chose spectrally close and bright, stable dyes (ATTO 633, 643, 655, 680, 700) to be evaluated as potential multicolor SD-DNA-PAINT candidates. A detailed description of the dye selection is outlined in Supplementary Information (Figure S1, Text S1). In short, the evaluation criteria for the dyes included their spectral property (emission spectra at 647 nm excitation) and the final image quality (brightness, background).

Based on these investigations we chose ATTO 655, ATTO 680 and ATTO 700 as promising dye candidates for triple-color SD-DNA-PAINT. To allow speed-optimized and simultaneous imaging of three distinct dye-imager conjugates (‘imagers’) within the same buffer, we chose three concatenated docking sequences (R3, R4, R6) that were recently introduced ^9^. We then designed three compatible imager sequences with matching melting points and a short spacer between dye and hybridization site to avoid guanine quenching of the ATTO dyes (R3S, R4S, R6S, Figure 2a). To evaluate the image quality in SD-mode, we first used these optimized imagers to perform single-color DNA-PAINT experiments in SD-mode on COS-7 cells immunolabeled for α-tubulin or clathrin (Figure 2b). Image processing was done as described below (Figure S2a, without color-filtering). The nanostructures of microtubules and clathrin coated vesicles were well resolved.

**Figure 2.**
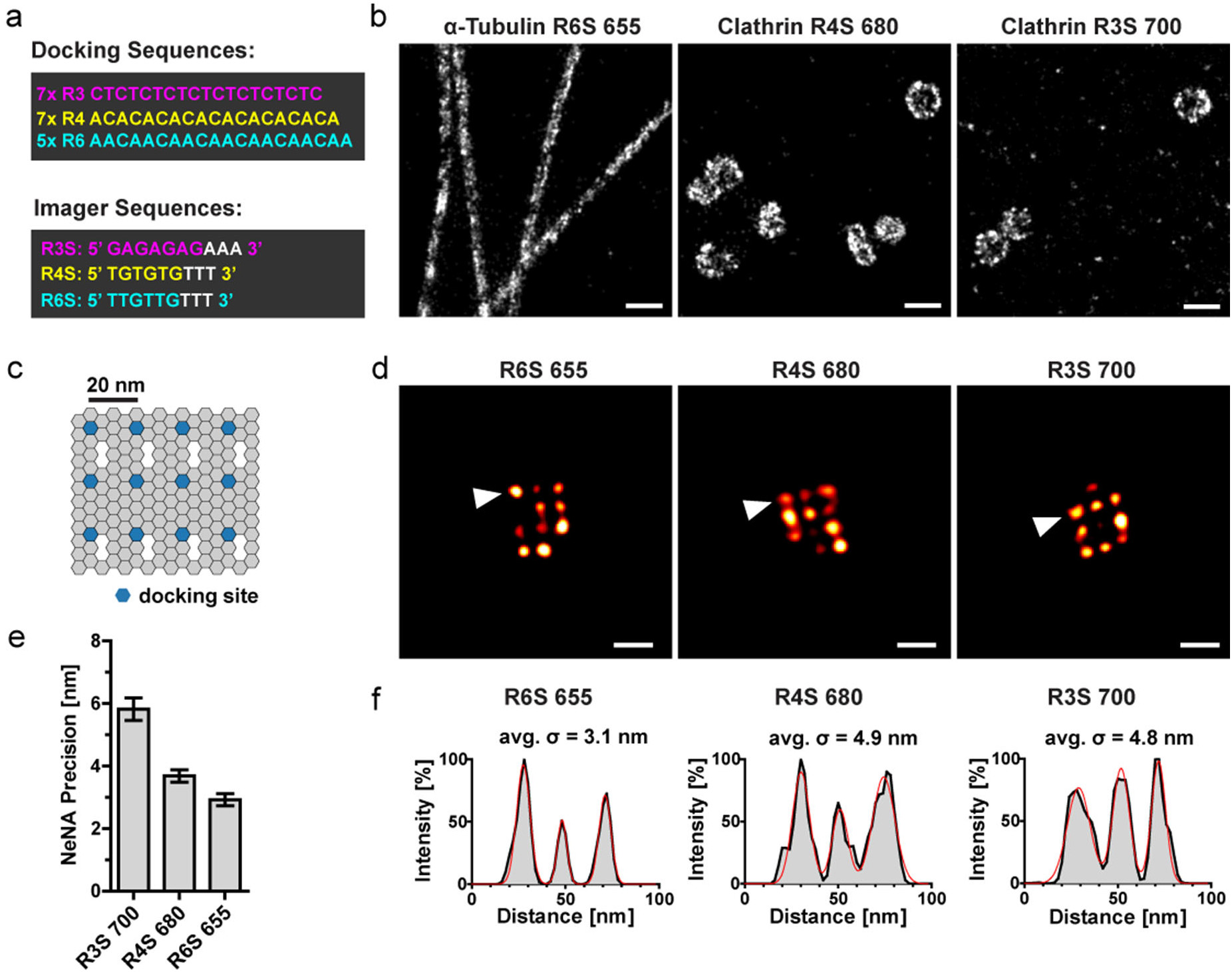
Image quality and localization precision. (**a**) Schematic of the oligonucleotide sequences of the docking strands (R3, R4, R6) and the optimized imager strands (R3S, R4S, R6S). (**b**) COS-7 cells where immunolabeled with primary antibodies against nanostructures (microtubules and clathrin coated vesicles) and secondary nanobodies conjugated to unique docking strands. Samples were imaged in SD-mode without color-filtering, using the imagers coupled to the ATTO dyes (R6S 655, R4S 680, R3S 700) as indicated. Recording modality: 20,000 frames for ATTO 680; 30,000 frames for ATTO (655/700), 100 ms exposure, 1 nM imager concentration. Selected regions show super-resolved nanostructures. Scale bar: 200 nm. (**c**) Schematic of the DNA origami structure with a 3 × 4 docking site arrangement with 20 nm grid distances. (**d-f**) Immobilized 20 nm DNA origamis with the docking sequences (R3, R4, R6) were imaged in SD-mode without color filtering, using the optimized imager strands (R3S, R4S, R6S) coupled to the ATTO dyes (655, 680, 700) as indicated. Recording modality: 20,000 frames, 100 ms exposure, 1nM imager. (**d**) Representative SD-DNA-PAINT images of DNA origamis. Scale bar: 40 nm. (**e**) The NeNA precision was calculated on the same single-color dataset as the images in (**d**). Mean +/− SEM, Images: 655 (n = 5), 680 (n = 4), 700 (n = 3). (**f**) Line profiles through the origamis (arrowheads in (**d**)) were fitted with a multi-Gaussian distribution (red line). The standard deviation of each single Gaussian was averaged (avg. σ).

To determine the localization precision and to clearly demonstrate the nanoscale resolution we next performed single-color DNA-PAINT experiments using the optimized imagers on 20 nm DNA origami grids (Figure 2c). The imaging was done in SD-mode and the data was processed as described below (Figure S2a, without color-filtering). The resulting images clearly show the resolved 20 nm patterns of the origami structures (Figure 2d). Next, we determined the experimental localization precision by performing a nearest neighbor analysis on subsequent localizations (NeNA precision) ^17, 18^. The values were between 3 and 6 nm for all used ATTO dyes (Figure 2e), similar to previously achieved precisions using Cy3b-labeled imagers in Exchange-PAINT (5 nm, ^9^). To verify this precision, we performed intensity line scans through the origami spots. The standard deviation (σ) of the Gaussian fits along the line scans were between 3.1 and 4.8 nm, demonstrating that the NeNA precision (Figure 2e) is an accurate estimate for the experimentally achieved localization precision (Figure 2f).

To perform dual-color SD-DNA-PAINT we chose the ATTO 655/700-imagers as those showed spectrally clearly distinct intensity populations (Figure S1c, e). To design color separation masks we performed single-color SD-DNA-PAINT experiments on COS-7 cells immunolabeled for abundant cellular structures (microtubules, clathrin). We used secondary nanobodies coupled to the described docking strands (R3, R6) and acquired the data in SD-mode using the corresponding (optimized) imagers (R6S-ATTO 655, R3S-ATTO 700). The color separation masks were designed to minimize the channel crosstalk while maximizing the inclusion of localizations (Figure 3a). For that, we quantified the crosstalk from the R6S-ATTO 655 channel (source) into R3S-ATTO 700 channel (recipient) and *vice versa* based on the single-color ground truth data. The masks were designed based on merged single-color intensity histograms to tolerate a maximal color crosstalk of 2% between both channels (Figure 3b). These masks included more than 70% of all localizations for the reconstruction of the images (Figure S3a).

**Figure 3.**
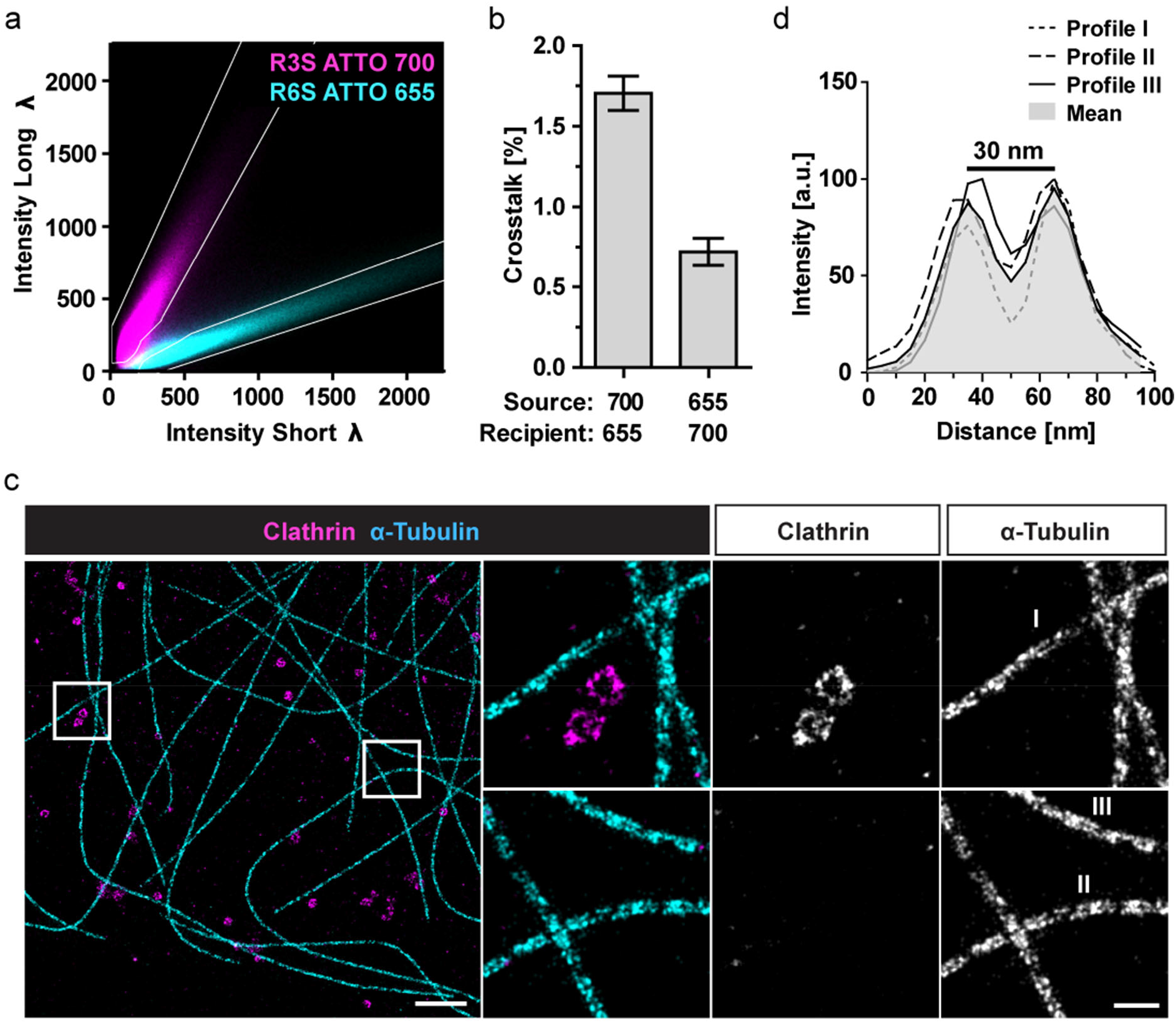
Dual-color SD-DNA-PAINT. **(a, b)** Single-color DNA-PAINT experiments in SD-mode using COS-7 cells immunolabeled for either clathrin (R3) or α-tubulin (R6). Recording modality: 20,000 frames, 100 ms exposure, 1 nM imager concentration. **(a)** Pseudo color overlay of the 2D intensity histograms from separate single-color experiments (cyan: R6S-ATTO 655, magenta: R3S-ATTO 700, both: average of 6 experiments). Color-separation masks (white lines) were manually designed within the 2D intensity histogram based on the single-color experiments. **(b)** The crosstalk was calculated as the percentage of localizations from the source (i.e. ATTO 700) that were detected within the mask of the recipient channel (i.e. ATTO 655) based on the single-color ground truth, and vice versa. Mean ± SEM, N = 4 – 5 images per channel. **(c)** COS-7 cells where immunolabeled for clathrin (R3) or α-tubulin (R6) simultaneously. Samples were imaged simultaneously in SD-mode (including color-filtering) using the imagers from (a). Representative dual-color SD-DNA-PAINT image of microtubules (R6S 655) and clathrin (R3S 700) with enlarged regions (white boxes) are shown. Scale bar: 1 μm, zoom 200 nm. Both enlarged regions have the same contrast. Recording modality: 30,000 frames, 100 ms exposure, 1 nM total imager concentration. **(d)** Intensity line profiles across the indicated microtubules shown in (c). The mean profile shows a ‘valley-to-peak’ intensity ratio of ~ 50% (dotted red line).

Next, we performed a dual-color SD-DNA-PAINT experiment on double immunolabeled COS-7 cells, stained for microtubules and clathrin with both imagers (R6S-ATTO 655, R3S-ATTO 700) in the buffer (Figure 3c). The data was processed using the optimized color separation masks as described above (Figure S2a). The resulting dual-color SD-DNA-PAINT images show microtubules and clathrin structures clearly separated with a very low crosstalk. The nanoscale structures of both tubular microtubules (double line) and spherical clathrin coated vesicles (ring structure) are clearly resolved. The peak-to-peak distance of the intensity line profiles across the microtubules (30 nm, Figure 3d) is in agreement with the distances ( ~ 29 nm) that have previously been calculated for similarly labeled microtubules ^19^.

The steps used to process the imaging data for dual-color SD-DNA-PAINT include the localization of single-molecule emissions, the drift correction, the pairing and grouping of the localizations, the color-filtering and the rendering of the multicolor image (Figure S2a). All those have been implemented in similar ways in previous SD-dSTORM approaches ^13, 14^. Here, we would like to discuss the color-filtering step in more detail and highlight the implementation of a novel multichannel registration procedure to further improve the precision and image quality of any SD-based approach, including SD-DNA-PAINT or SD-dSTORM.

One seemingly critical point of the SD-mode was the exclusion of localizations through the color-filtering using the color separation masks ^12^. In dual-color SD-DNA-PAINT, for example, about 30 % (Figure S3a) of the initial localizations plotted within the 2D intensity histogram (Figure 3a) are excluded by the masks to keep the crosstalk below 2% (Figure 3b). This exclusion might be conceived as a disadvantage. However, most of the excluded localizations in fact had low intensity values (Figure 3a) and were therefore less precise compared to the included localizations with high intensities. To demonstrate that, we reconstructed images from cellular nanostructures (Figure S2b) and 20 nm DNA origami grids (Figure S2c) using the included or excluded localizations of several dyes. While for some conditions, structures were still partially visible in the images rendered from the excluded localizations, these images were of much lower quality and contained grossly deteriorated nanostructures in comparison to the correctly filtered (‘included’) images. Therefore, the applied color filtering procedure reduces the number of low-quality localizations and thus enhances the quality of the super-resolved images.

In multicolor SMLM the accuracy of color channel registration is of central importance for any experiments in which relative nanoscale distances between different objects are measured. In SD-mode even mild flat-field distortions within the optical path (e.g. emission splitter) could contribute significantly to an error in registration of the short and long wavelength channels. Originally, potential errors in multichannel registration were prevented by reconstructing the final image using only the single-molecule localizations of one (i.e. short) wavelength channel ^12, 13^. This resulted in a partial loss of the single-molecule emissions that were split into the long wavelength channel. Here, we optimized this approach by harvesting the single-molecule emissions from both channels to achieve the highest possible localization accuracy. To correct for nanoscale distortions between the split channels we designed a custom registration and unwarping procedure that was implemented after the color-filtering step (Figure S2a). In this procedure we used the entire population of paired localizations as intrinsic fiducial markers for correction of the local offset between the split channels.

In addition to this registration procedure, we used the intensity values of the localization in the short and long wavelength channels to calculate an intensity-weighted localization of each localization pair. This intensity-weighted localization correction ensured that we take advantage of the entire detected emission from each single-molecule event to determine its precise localization. Compared to using only one of the channels (i.e. short) this novel intensity-weighted multichannel registration procedure significantly improved the image quality on cellular nanostructures (Figure. S2b) and 20 nm DNA origami grids (Figure S2c). Importantly, the localization precision measured from DNA origamis was significantly improved using this procedure. For ATTO 700 that has the weakest intensities in the short wavelength channel, the NeNA precision was almost doubled from 11 nm to 6 nm (Figure S2c). Importantly, this intensity-weighted multichannel registration procedure uses the localizations of the structure under investigation as fiducial markers rather than exogenously added fiducials (e.g. beads, nanogold). That way, the structural signal governs the corrections rather than the signal from regions that are normally excluded from the final image.

As discussed earlier, DNA-PAINT achieves higher localization precisions compared to dSTORM approaches, because it offers a higher signal-to-noise ratio for the single-molecule emissions. To assess the difference in performance between dSTORM and DNA-PAINT using SD-mode, we directly compared the recently established multicolor SD-dSTORM ^12, 13^ with SD-DNA-PAINT. Since SD-dSTORM was so far limited to two colors, we directly compared it to dual-color SD-DNA-PAINT using the same microscope system. For SD-dSTORM, the emission splitter was equipped with a distinct dichroic mirror for optimal color separation of the specific dyes. Again, we used COS-7 cells immunolabeled for microtubules and clathrin with method-specific secondary antibodies. For direct comparison of the image quality, we used the same laser power settings and image processing steps as described above for SD-DNA-PAINT (Figure S2a). SD-DNA-PAINT achieved a significantly higher localization precision compared to SD-dSTORM (Figure 2e, Figure S4c). In line with that, the cellular nanostructures in the SD-dSTORM images (double line of microtubules, ring structure of clathrin coated vesicles, Figure S4a) were not as clearly resolved as in the SD-DNA-PAINT images (Figure 3c). The intensity line profiles across the microtubules showed more prominent ‘valley-to-peak’ ratios for SD-DNA-PAINT (Figure 3d) compared to SD-dSTORM (Figure S4b). Similar to SD-DNA-PAINT, SD-dSTORM also gains precision by implementing the novel intensity-weighting procedure (Figure S4c).

Next, we aimed at performing triple-color SD-DNA-PAINT using the spectrally suited triplet of ATTO dyes (655/680/700) conjugated to distinct imagers (R6S, R4S, R3S). To optimize the color separation masks for triple-color we first performed single-color SD-DNA-PAINT experiments as described above. The emission spectrum of ATTO 680 peaks in between the spectra of ATTO 655 and ATTO 700 (Figure S1a, b). The 2D intensity histogram of the overlaid single-color experiments clearly shows three distinct populations of localizations with an overlap of ATTO 680 with ATTO 655 and ATTO 700 at low intensities (Figure 4a; Figure S1c-e). In analogy to the dual-color masks, the triple-color separation masks were designed to minimize color crosstalk while maximizing the inclusion of localizations. Despite the spectral overlap of ATTO 680 with both ATTO 655 and ATTO 700, up to 70% of the localizations could be included (Figure S3c) while tolerating a maximal crosstalk of 5% (Figure 4b). After optimizing the triple-color separation masks, we performed triple-color SD-DNA-PAINT on cells immunolabeled for three cellular nanostructures (microtubules, clathrin coated vesicles, vimentin filaments) with all three imagers (R6S-ATTO 655, R4S-ATTO 680, R3S-ATTO 700) in the imaging buffer. The images were processed as described above using the optimized triple-color separation masks. The resulting SD-DNA-PAINT images demonstrate that the nanoscale structures of all three biological structures are clearly resolved with minimal crosstalk using the triple dye combination ATTO 655/680/700 (Figure 4c). The low crosstalk in triple-color SD-DNA-PAINT was verified using DNA origami grids with distinct docking strands (Figure S3c).

**Figure 4.**
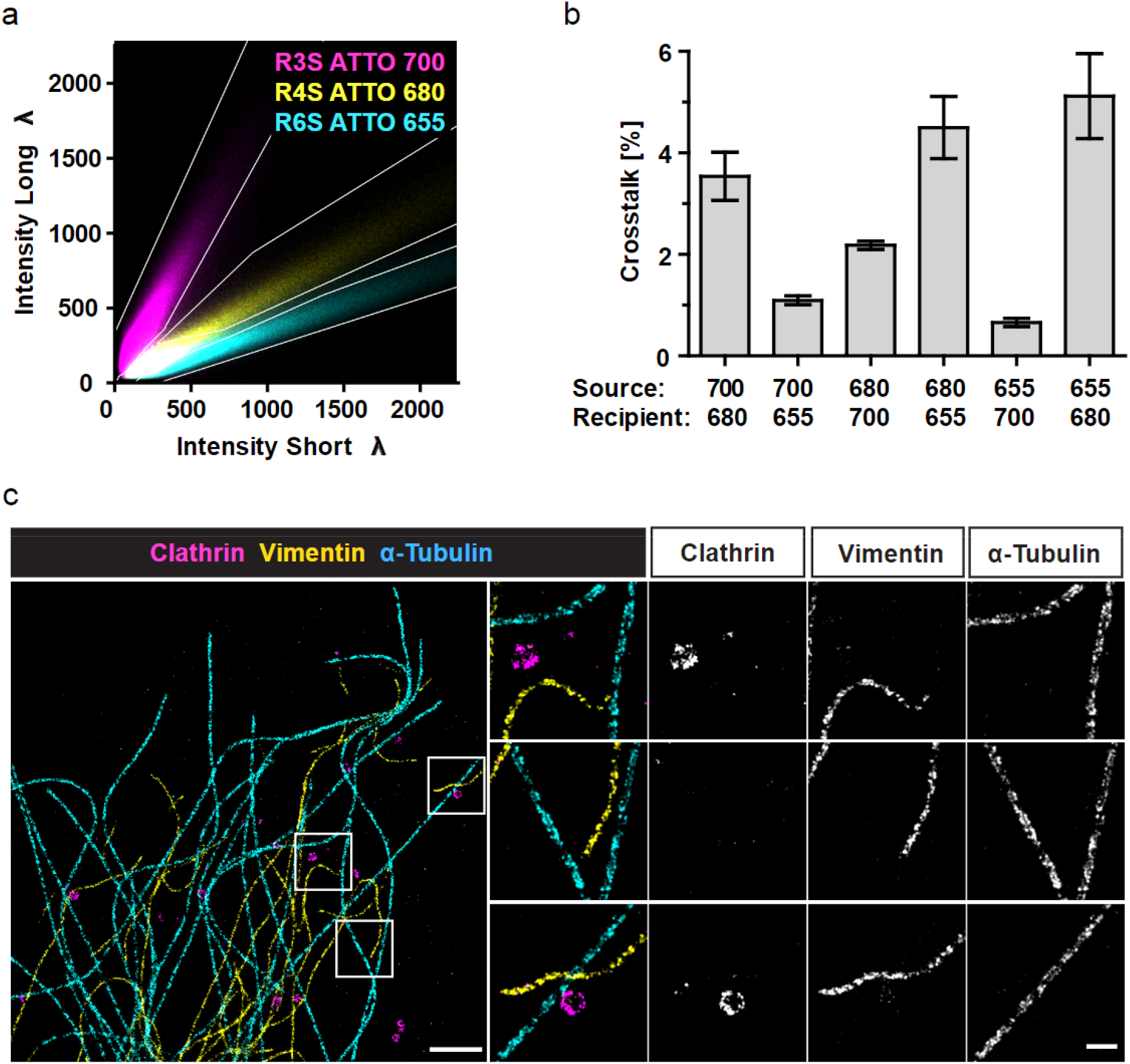
Triple-color SD-DNA-PAINT. (**a,b**) Single-color DNA-PAINT experiments were performed in SD-mode using COS-7 cells immunolabeled either for clathrin (R3S, R4S) or α-tubulin (R6S). Recording modality: 20,000 frames, 100 ms exposure, 1 nM imager concentration. (**a**) Pseudo-color overlay of the 2D intensity histograms from separate single-color experiments (cyan/magenta/yellow: R6S 655 / R3S 700 / R4S 680). Color-separation masks (white lines) were manually designed based on the single-color signals to minimize the crosstalk between the channels while maximizing the included localizations. Histograms from 5 - 6 images were averaged per channel. (**b**) The crosstalk was calculated for all dye combinations as the percentage of localizations from the source that are detected within the mask of the recipient channel based on the single-color ground truth. The triple-color masks were designed to tolerate a crosstalk below 5 %. Mean ± SEM, N = 4 – 5 images per channel. Data in (**a, b**) was partially reused from Figure 3. (**c**) COS-7 cells immunolabeled for clathrin, vimentin and α-tubulin were imaged in SD-mode, using the imagers from (a). Representative triple-color SD-DNA-PAINT image and indicated sub-regions (white boxes) of microtubules (R6S 655), vimentin (R4S 680) and clathrin (R3S 700) are shown. Scale bar: 1 μm, zoom 200 nm. All three regions have the same contrast. Recording modality: 30,000 frames, 100 ms exposure, 1 nM total imager concentration.

Compared to the original SD-dSTORM approach ^12^ our novel intensity-weighted multichannel registration procedure is not intrinsically free from chromatic distortions, instead, these errors are corrected. To verify the high accuracy of this procedure in triple-color SD-DNA-PAINT mode we imaged 20 nm DNA origami grids containing a single docking strand (R6) with three compatible imagers (R6S-ATTO 655/680/700) in the imaging buffer (Figure S5a). If the registration is precise, all three color channels should show a high degree of overlap. Indeed, the resulting triple-color overlay shows a clear colocalization of the docking site spots in all three channels. To quantify the accuracy of multichannel registration, we performed a nearest neighbor (NN) distance analysis between each of the channels on the origami data set (Figure S5c, d). The peaks of the multichannel NN distance distributions were significantly above the control for a random distribution and in the range of 1 – 3 nm for all comparative color channel combinations. This demonstrates the high accuracy of our custom-designed multichannel registration procedure that corrects for flat-field distortions of the emission splitter with an error that is below the localization precision of single-color images.

## CONCLUSIONS

In summary, we presented a novel approach that combines DNA-PAINT with simultaneous multichannel acquisition using spectral demixing (SD). SD-DNA-PAINT alleviates the need for sequential fluid exchange and therefore avoids all associated disadvantages, including the extended acquisition times. SD-DNA-PAINT speeds up the acquisition time of current multichannel DNA-PAINT while taking advantage of all the benefits of DNA-PAINT. To realize SD-DNA-PAINT we selected a novel combination of spectrally overlapping dyes (ATTO 655/680/700) and designed DNA-PAINT probes for optimal color separation and resolution. Using these probes, we achieved simultaneous multicolor DNA-PAINT with minimal crosstalk and excellent localization precision (3 – 6 nm, Figure 2e). We demonstrated the ability of SD-DNA-PAINT to resolve 20 nm DNA origami grids and cellular nanostructures.

The presented SD-DNA-PAINT approach offers up to three color channels. Including more colors was limited by the partial overlap of the dye-specific intensity populations within the histogram that is used for color separation (Figure 4a). The laser power density of the SMLM system used here was limited to max. 3 kW/cm^2^, while other applications (e.g. ultra-high resolution DNA-PAINT ^8^) used higher power densities (> 4 kW/cm^2^). We propose that by using higher laser powers the intensity populations within the histogram could be shifted towards higher values and thus create sufficient space for additional dye channels.

A large number of researchers, including ourselves, routinely used variants of multicolor dSTORM to resolve cellular nanostructures ^13, 15, 20–22^. Compared to SD-dSTORM, SD-DNA-PAINT offers higher localization precisions and a third color channel. In analogy to qPAINT ^7^, the controlled binding kinetics of SD-DNA-PAINT could be used to count binding sites in biological structures. We therefore believe that SD-DNA-PAINT will become a useful method to perform robust, fast and quantitative multicolor SMLM in cases where simultaneous acquisition of up to three color channels are desired.

## ABBREVIATIONS

SMLM: Single-molecule Localization Microscopy
STORM: Stochastic Optical Reconstruction Microscopy
SD: Spectral Demixing
PAINT: Points Accumulation in Nanoscale Topography
fm-DNA-PAINT: Frequency Multiplexing DNA-PAINT
NeNA: Nearest Neighbor Based Analysis
NN: Nearest Neighbor
AB: Antibody
NB: Nanobody
F(ab’)1: Antigen Binding Fragment

## ASSOCIATED CONTENT

### Supporting text

Detailed experimental methods and material; Supporting Text to S1: Dye selection for SD-DNA-PAINT.

### Supporting figures

Supporting Figure S1: Dye Selection; Supporting Figure S2: Image Processing; Supporting Figure S3: Color filtering; Supporting Figure S4: SD-dSTORM; Supporting Figure S5: Multichannel SD-DNA-PAINT registration accuracy.

### Supplementary Tables

Supplementary Table 1: Imager Sequences; Supplementary Table 2: Docking Sequences; Supplementary Table 3: Experimental settings.

### Supplementary Data

SD-DNA-PAINT filter set (file type: PNG); SD-dSTORM filter set (file type: PNG); SDmixer configuration file (file type: TXT); ThunderSTORM configuration file for SD-DNA-PAINT (file type: TXT); ThunderSTORM configuration file for SD-dSTORM (file type: TXT).

## AUTHOR INFORMATION

### Author Contributions

J.S. and N.G. conceived the scientific concept and designed the experiments. N.G. performed the experiments, designed the software tools and analyzed the data. J.S. supervised the study and together with N.G. wrote the manuscript. S.S. and R.J. contributed DNA-PAINT reagents and expertise in how to perform experiments. R.J. supervised S.S. and gave advice on structure and content of the manuscript. All authors contributed to writing of the manuscript.

### Note

The authors declare no competing financial interest.

### Funding Sources

This work has been supported in part by the German Research Foundation (DFG) through the SFB958 (project Z02) to J.S., and in part by the European Research Council through an ERC Starting Grant (MolMap; grant agreement number 680241) and the Max Planck Society.

## ACKNOWLEDGEMENT

We thank the staff of the Advanced Medical Bioimaging Core Facility, Charité-Universitätsmedizin, Berlin, Germany (AMBIO) for support in sample preparation and usage of microscope systems. We thank Dr. Hylkje Geertsema (TU Delft) from the laboratory of Prof. Helge Ewers (FU Berlin) for initial discussions about DNA-PAINT probes.

## Notes

https://github.com/gtadeus/sdmixer2

https://github.com/ngimber/Converter_ThunderSTORM_SDmixer

https://github.com/ngimber/SD-DNA-PAINT

https://github.com/ngimber/SMLM_VoronoiTesselation

https://github.com/ngimber/SMLM_NearestNeighbor

### Competing Interest Statement

The authors have declared no competing interest.

