## Supplementary Information for "Simultaneous multicolor DNA-PAINT without sequential fluid exchange using spectral demixing"

**Supporting Figure S1.** Dye Selection.

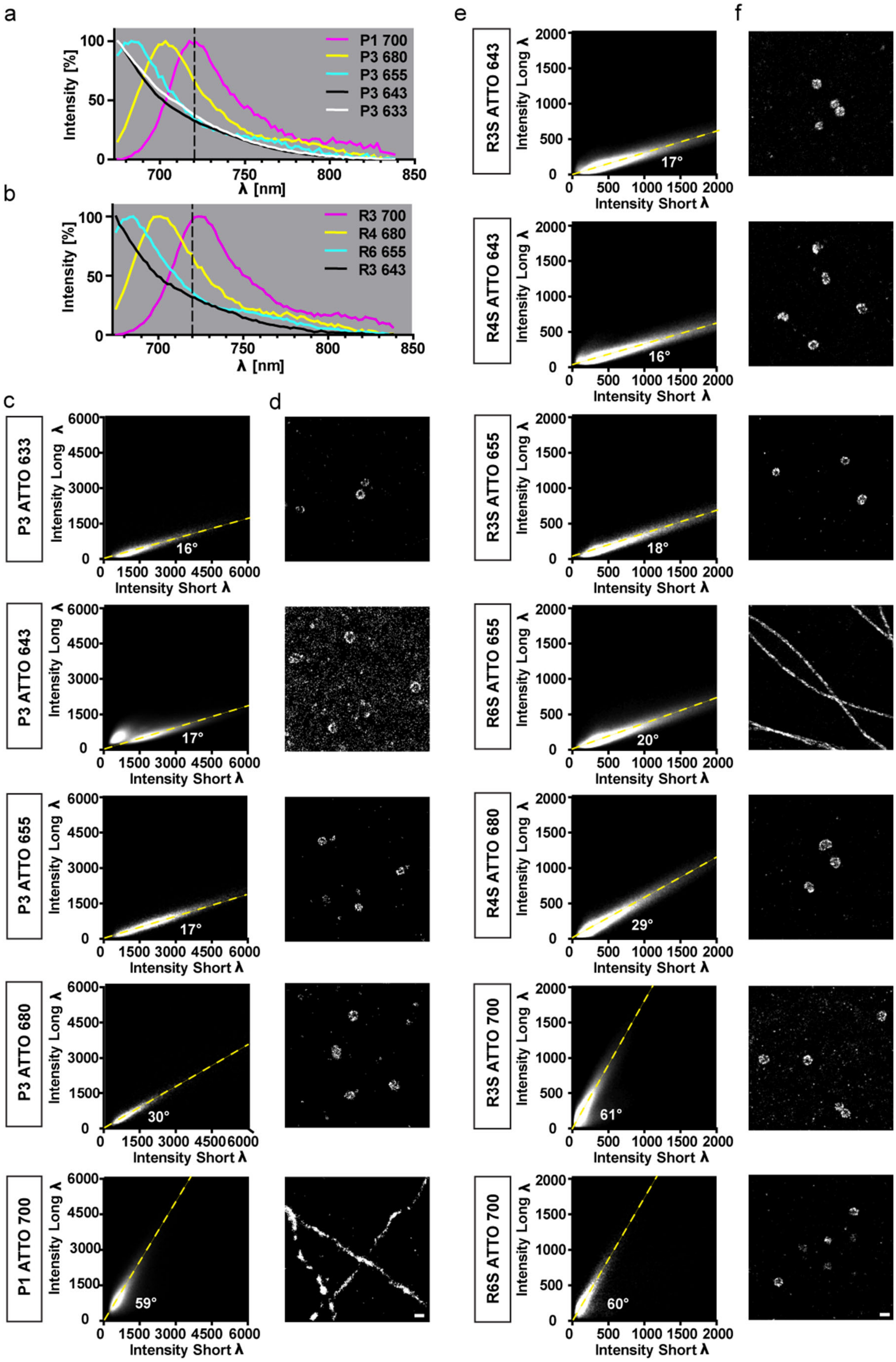

**Supporting Figure S1.** Dye Selection. (**a, b**) Normalized emission spectra of the used imagers (**a**) P1 / P3 and (**b**) R3S / R4S / R6S. The dashed lines indicate the position of the beam splitter. (**c - f**) Single-color DNA-PAINT experiments were performed in SD-mode on COS-7 cells immunolabeled either for clathrin (P3, R3S, R4S) or α-tubulin (P1, R6S). Recording modality: 20,000 frames, 100 ms exposure, 1 nM imager concentration. The angles between the major direction of propagation (yellow line) and the x-axis are indicated. (**c, d**) 2D intensity histograms (**c**) and example images (d) of the candidate imagers P3 ATTO 633 (N = 5), P3 ATTO 643 (N = 5), P3 ATTO 655 (N = 6), P3 ATTO 680 (N = 4) and P1 ATTO 700 (N = 6). Fixation method: Methanol. (**e, f**) 2D intensity histograms (**e**) and example images (**f**) of the candidate imagers R3S ATTO 643 (N = 5), R4S ATTO 643 (N = 3), R3S ATTO 655 (N = 3), R6S ATTO 655 (N = 6), R4S ATTO 680 (N = 5), R3S ATTO 700 (N = 6), R6S ATTO 700 (N = 4). Fixation method: Glutaraldehyde.

### Supporting Text S1: Dye selection for SD-DNA-PAINT

Multicolor SD-DNA-PAINT required dyes with sufficiently overlapping emission to be excited by the same laser line. At the same time, the spectral shift between the dyes had to be sufficiently large to distinguish the dyes in SD-mode according to their intensities in the short and long wavelength channel. In addition, high brightness was desired to achieve a high localization accuracy. The third criterion was the hydrophobicity of the dye, which, if too high, can cause unwanted background localizations through unspecific binding of the dye-imager strand to the substratum. Five dye candidates that fulfilled all mentioned criteria were the spectrally close ATTO dyes (633, 643, 655, 680, 700) which we evaluated according to the properties of the 2D intensity histogram and the resulting image quality (brightness, background).

For an initial selection, we coupled the standard imagers P1, P3 to each dye candidate [18]. When excited at 647 nm, the emission spectra of those imagers showed closely overlapping emission spectra (Figure S1a). Clearly, the emission spectra of the ATTO 633, 643, 655 imagers were too similar to be separated using spectral demixing. The main emission peaks of ATTO 680 and 700, in contrast, were both shifted to longer wavelengths and therefore remained candidates for triple-color SD-DNA-PAINT in combination with one of the other imagers. To evaluate the image quality and the feasibility of separating the imagers using SD-mode we performed single-color SD-DNA-PAINT experiments on COS-7 cells that were immunolabeled for abundant cellular proteins (α-tubulin, clathrin) that reveal characteristic nanostructures in high quality super-resolution images (Figure S1d). Image data was processed as described (Figure S2a). To allow good separation of the probes the 2D intensity distributions should exhibit high values and should be as tight as possible. The image quality, including unwanted background, was the other criteria for selection. As expected from the emission spectra (Figure S1a), the 633/643/655-imagers showed strongly overlapping populations at similar angles within the 2D intensity histogram. In contrast, the 680/700 imagers showed distinct angles. We repeated the experiments with the ATTO dyes 643, 655, 680, 700 coupled to the optimized imager sequences R3S, R4S, R6S on COS-7 cells fixed in glutaraldehyde. The images using the imagers revealed different levels of background. ATTO 633/655/680/700 had moderate to low background after fixation with glutaraldehyde or methanol, while ATTO 643 showed a high, unspecific background signal specifically after methanol fixation (Figure S1d,e). In particular, the 655 population showed both very high intensity values and a tight intensity distribution. We therefore used the dye combination ATTO 655/680/700 for all other experiments.

**Supporting Figure S2.** Image Processing.

a

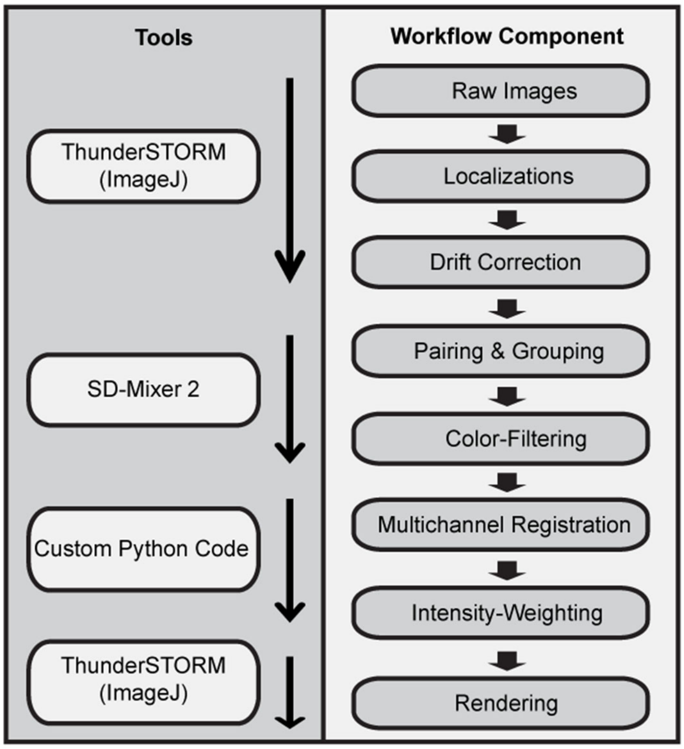

b

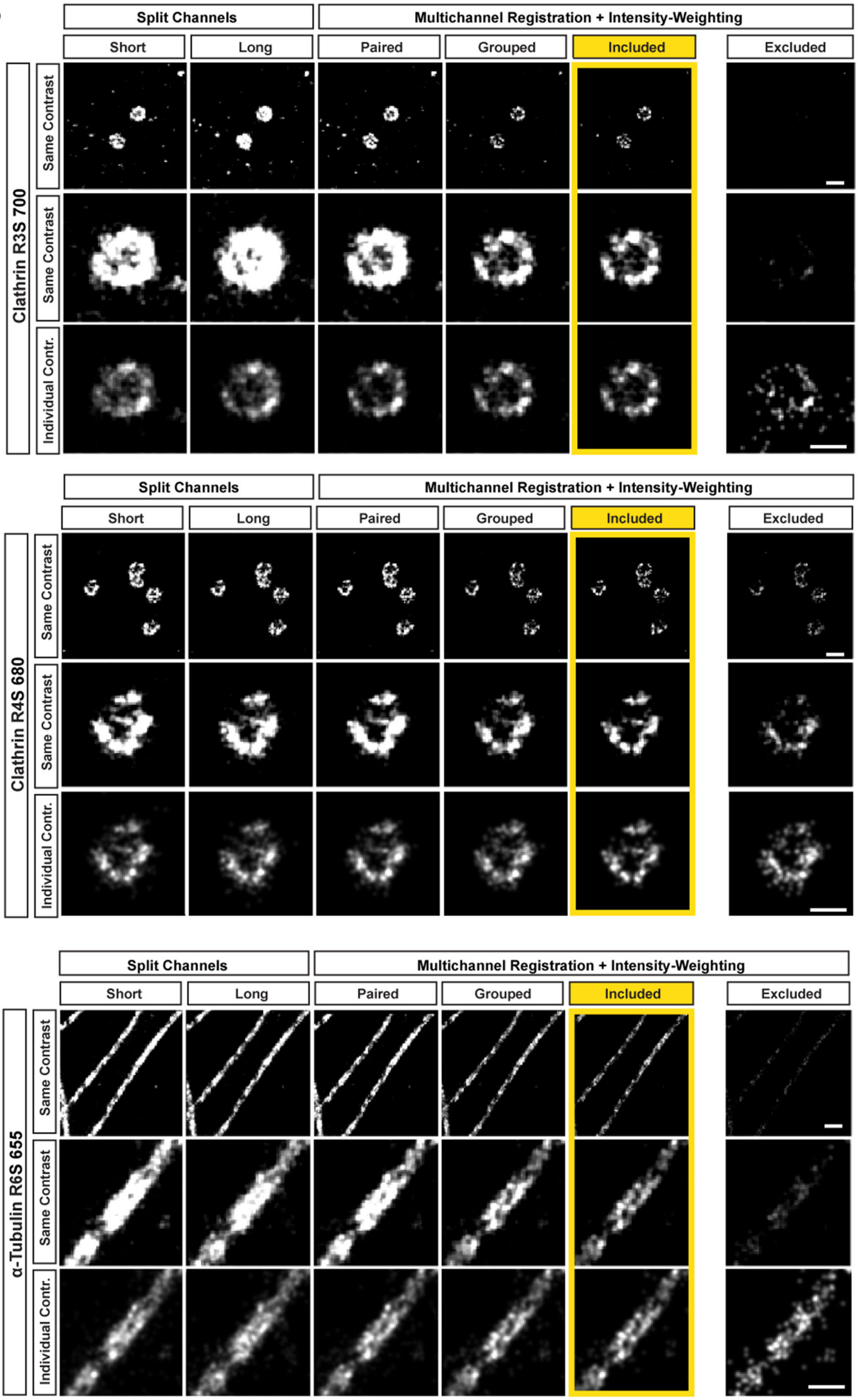
Split Channels
Multichannel Registration + Intensity-Weighting
Short
Long
Paired
Grouped
Included
Excluded
Clathrin R3S 700
Same Contrast
Same Contrast
Individual Contr.
Clathrin R4S 680
α-Tubulin R6S 655

c

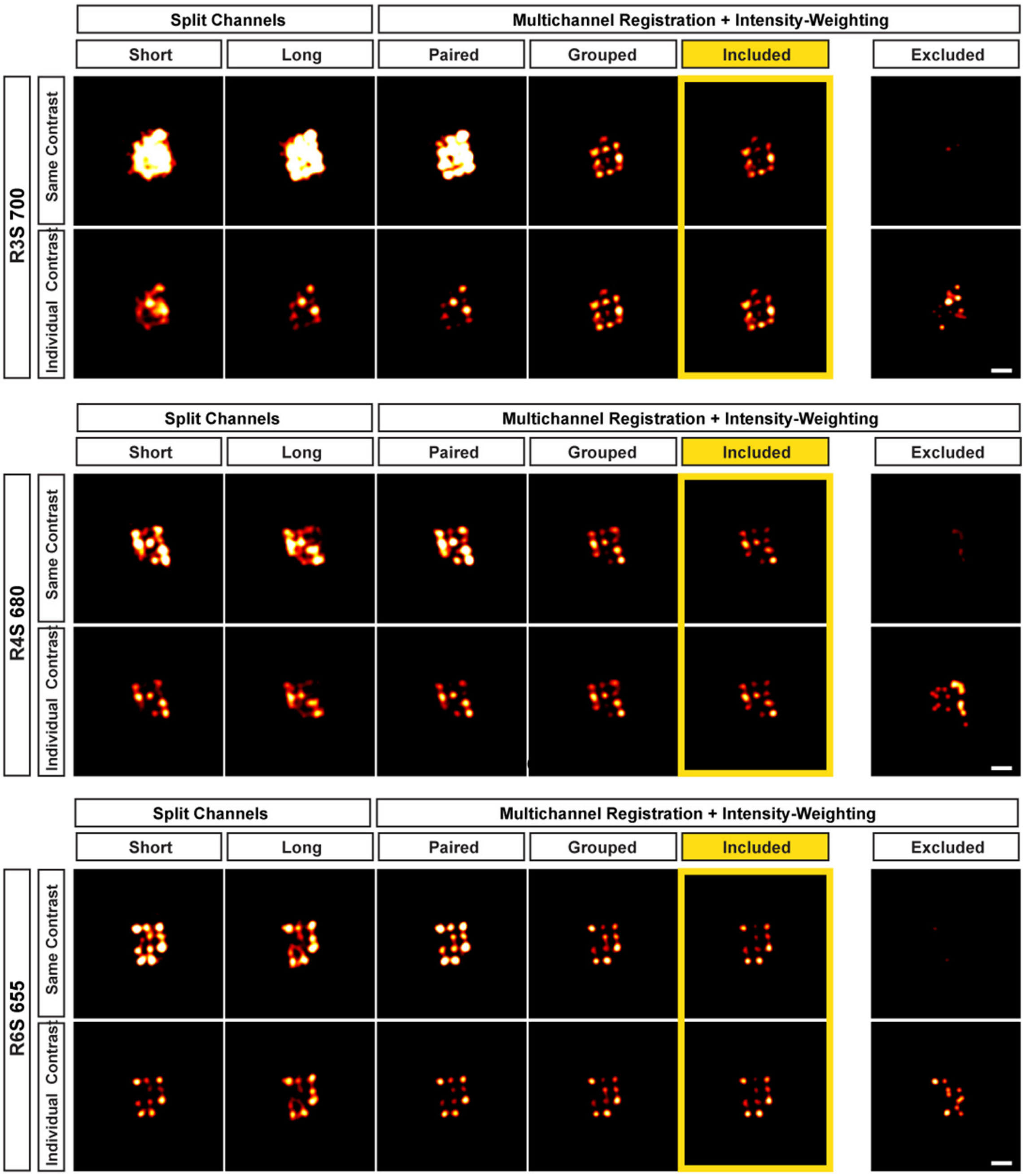
Split Channels
Multichannel Registration + Intensity-Weighting
Short
Long
Paired
Grouped
Included
Excluded
R3S 700
Same Contrast
Individual Contrast
R4S 680
R6S 655

d

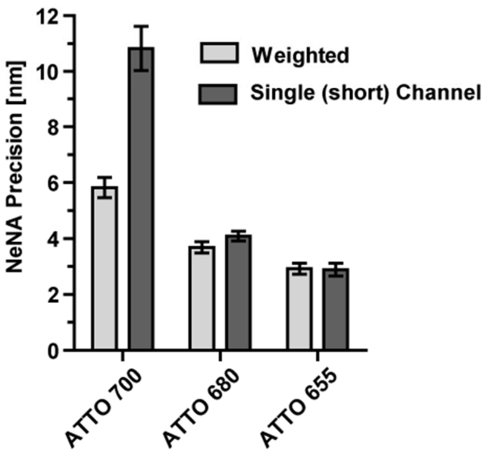
NeNA Precision [nm]
0
2
4
6
8
10
12
Weighted
Single (short) Channel
ATTO 700
ATTO 680
ATTO 655

**Supporting Figure S2**. Image Processing. (**a**) Schematic of the image processing workflow for SD-DNA-PAINT. Left column indicates the software tools and the right column indicates specific workflow component. (**b, c**) To compare the image quality at different stages of image processing within the SD-DNA-PAINT workflow, we performed SD-DNA-PAINT experiments on single-color samples using the indicated imagers (R3S 700, R4S 680, R6S 655) either on (**b**) immunolabeled COS-7 cells (clathrin or α-tubulin) or (**c**) on 20 nm gridded DNA origamis. Recording modality: 20,000 frames for origamis and clathrin 680; 30,000 frames for clathrin 700 and α-tubulin 655; all 100 ms exposure, 1 nM imager concentration. Example images (top rows) and regions of interest (middle rows) are shown at different processing steps, with the same contrast, to demonstrate changes in intensity levels. Adjusting the contrast individually, to visualize low signals (bottom rows), uncovered structural differences. The processing steps indicated are the following: 'Short' / 'Long' = Image of separate channels on the camera before pairing; All other processing steps include multichannel registration and intensity-weighting: 'Paired' = Image after pairing the localizations; 'Grouped' = Image after grouping the paired localizations; 'Included' (yellow framing, final image) = Image rendered from the grouped localizations included in the masks; 'Excluded' = Image rendered from only the grouped localizations that were excluded from the masks. Note the increase in image quality after pairing and intensity-weighting and that the signal from the excluded localizations is very weak compared to the included localizations (yellow frame). Nanostructures are barely visible in the excluded images, even after individual contrast enhancement. Scale bars: cells 100 nm, DNA origamis 40 nm. (**d**) The NeNA precision was calculated on the single-color data set used in (**c**) in SD-mode without grouping and color-filtering. Note the improved precision of ATTO 700 upon intensity-weighting the localizations from both channels. The data is partially replotted from Figure 2e. Mean +/- SEM, Images: 655 (n = 5), 680 (n = 4), 700 (n = 3).

**Supporting Figure S3.** Color filtering.

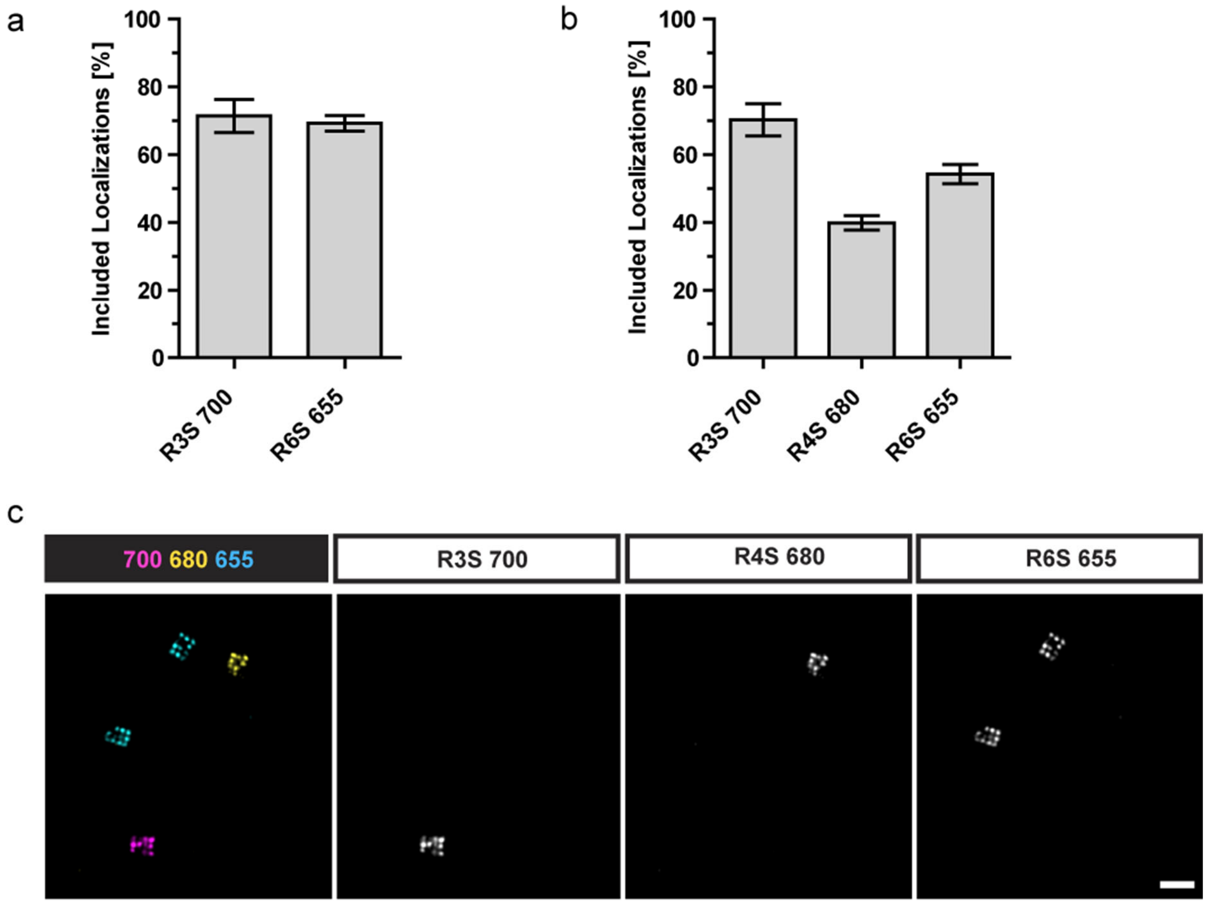


**Supporting Figure S3**. Color filtering. (**a, b**) To determine the proportion of excluded localizations during color filtering, dual and triple-color separation masks were applied to single-color DNA-PAINT experiments (in SD-mode) on COS-7 cells immunolabeled for clathrin (R6S-ATTO 655) or microtubules (R3S-ATTO 700). Recording modality: 20,000 frames, 100 ms exposure, 1 nM imager concentration. Quantification of the included localizations as a percentage of all grouped localizations for both dual-color (**a**) and triple-color (**b**) SD-DNA-PAINT experiments using the indicated imagers (R3S 700, R4S 680, R6S 655). Mean ± SEM, N = 4 – 5 images per channel. (**c**) To visualize the level of color channel crosstalk we performed a triple-color SD-DNA-PAINT experiment on a mix of immobilized DNA origami grids (20 nm spacing) each with distinct docking sequences (R3S magenta, R4S yellow, R6S cyan) using all three imagers (R3S 700, R4S 680, R6S 655) simultaneously. Scale bar: 100 nm. Recording modality: 50,000 frames, 100 ms exposure, 1 nM imager total.

**Supporting Figure S4.** SD-dSTORM.

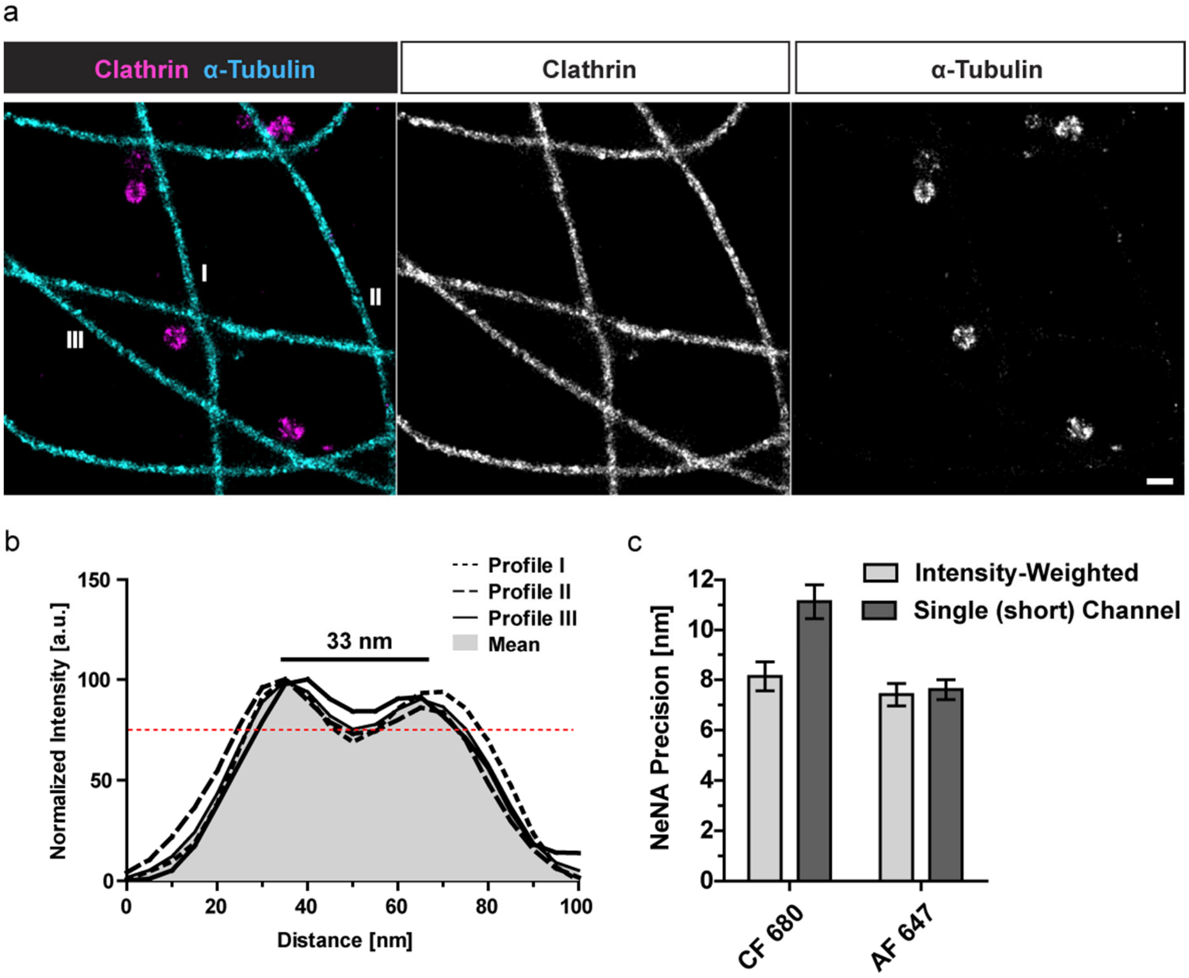


**Supporting Figure S4**. SD-dSTORM. (**a-c**) Dual-color SD-dSTORM experiments were performed on COS-7 cells immunolabeled (primary AB, secondary F(ab')1 fragment) for clathrin (CF 680) and microtubules (AF 647). The raw data was processed similar as described previously [13], but including the novel multicolor registration and intensity-weighting procedure (Figure S2). (**a**) The SD-dSTORM images show the nanoscale structures of clathrin coated vesicles (ring structure) and microtubules (double line profile). Scale bar: 200 nm. (**b**) Intensity line profile across microtubules at indicated regions in (a). The mean profile shows a 'valley-to-peak' intensity ratio of ~75% (dotted red line, compare to Fig. 3d). (**c**) NeNA precision of the same dataset using either the localizations from the short wavelength channel or weighted localizations. Note the improved precision of Al 647 upon weighting the intensities from both channels. Recording modality: 30,000 frames, 30 ms exposure.

**Supporting Figure S5.** Multichannel SD-DNA-PAINT registration accuracy.

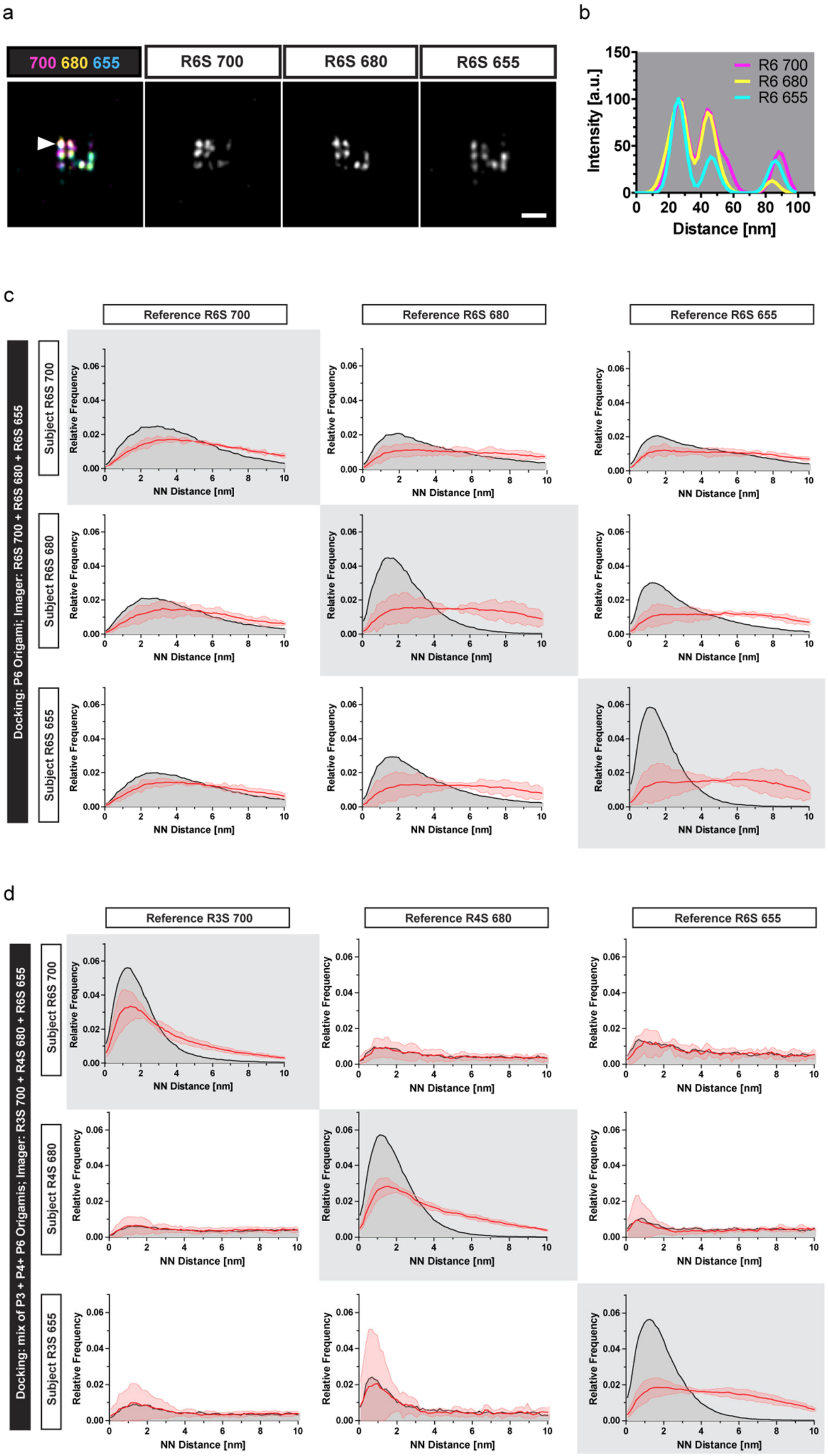

**Supporting Figure S5**. Multichannel SD-DNA-PAINT registration accuracy. **(a-c)** Triple-color SD-DNA-PAINT experiment were performed on a single type of 20 nm DNA origami grids (docking sequence R6) using the three imagers R6S 700, R6S 680, R6S 655 simultaneously. **(a)** Representative image. Scale bar: 50 nm. **(b)** Intensity line profiles through the origami (arrowhead in (a)). Note the precise overlap of all three color channels. **(c)** Nearest neighbor distance analysis (NN distance) between the channels serve as a measure for 'colocalization'. The graphs display the distance distributions of nearest 'reference' localization to each 'subject' localization. All possible reference/subject combinations are plotted as indicated. The grey-labeled graphs (diagonal from top left to bottom right) show the NN distance distributions for each channel to itself (positive control: 'colocalization'). The red lines indicate the upper and lower bounds of the 95% confidence interval of spatial randomness. All distributions in (c) show peaks above random between 1 and 3 nm, demonstrating that those channels are registered with an accuracy below the channel-specific localization precision (compare to Figure 2e). N = 3 images. **(d)** As a 'negative control' for the NN distance distributions the experiments were repeated with a mix of the 20 nm DNA origami grids R3, R4 and R6, imaged with R3S 700, R4S 680 and R6S 655 (see also Fig. S3c). Again, the NN distance distributions from each channel to itself (grey-labeled graphs, 'positive control') showed peaks above random. All other NN distance distributions ('negative controls') did not differ from random, demonstrating that the peaks in (c) originate from localizations at the same DNA origami docking site ('colocalization') and not from a random distribution (N = 5 images). Recording modality for all sub-figures: 50,000 frames, 100 ms exposure, 1 nM total imager concentration.

### METHODS

**Imager design:** We designed the three imager sequences (R3S, R4S, R6S, see also Table 1) to specifically bind the concatenated docking sequences R3, R4 and R6 [9] and to have similar off-rate kinetics for simultaneous multicolor acquisition. Therefore, the imagers were designed with comparable free energies of -12.7 kcal/mol (R3S), -11.5 kcal/mol (R4S), and 11.6 kcal/mol (R6S), calculated using NUPACK (600 mM NaCl and 20°C, [23]). The fluorophores ATTO 655, ATTO 680 and ATTO 700 are quenched by guanine bases, as known for oxazine fluorophores [24]. To minimize this effect, a few bases (adenine or thymine) were added as spacers to the imager sequences (Figure 2a). All imagers were purchased from Eurofins Genomics as a 3'-conjugation to the fluorophore. Fluorescence emission spectra from the imagers were validated with a spectro-fluorimeter (Varioskan Flash, Thermo Scientific) with 647 nm laser excitation.

**DNA origami folding and purification:** All DNA origami structures were designed using Picasso's 'Design' tool [18]. Folding and purification of DNA origami structures was performed as described previously [9].

**DNA Origami immobilization:** DNA Origami nanostructures were assembled as described above and immobilized on µ-Slide chambers (8-well #1.5, Ibidi). Chambers were washed with PBS, incubated with 200 µl biotin-labeled BSA (1 mg/ml, Sigma-Aldrich, A8549) in PBS, 5 min) and washed again. Chambers were then incubated with 200 µl neutravidin (1mg/ml [Thermo Scientific, 310000] in PBS, 5 min) and washed with PBS + 10 mM $MgCl_2$ ('$PBS^{+}$'). 3 µl DNA were diluted in 200 µl $PBS^{+}$ and applied to the chamber. The chambers were washed after 5 min with $PBS^{+}$. This step was repeated if the origami density was too low. 150 nm gold nanoparticles (Cytodiagnostics , G-150-20) were diluted 1:10 in $PBS^{+}$ and applied to the imaging chambers. Gold particles were allowed to settle onto the bottom (5 min RT) and washed with PBS$^{+}$. Immobilized nanostructures were then used for SD-DNA-PAINT experiments.

**DNA / Nanobody conjugation:** Nanobodies against rabbit IgG (10E10) and mouse kLC (clone 1A23) were purchased from Nanotag with a single ectopic cysteine at the C-terminus for site-specific and quantitative conjugation. Conjugation to DNA-PAINT docking sites (Table 2) was performed similarly to the previously described method [25]. First, buffer was exchanged to PBS + 5 mM EDTA, pH 7.2 using Amicon centrifugal filters (10k MWCO), then free cysteines were reacted with 20-fold molar excess of bifunctional DBCO-PEG$_4$-Maleimide linker (Jena Bioscience, CLK-A108P) for 2–3 h on ice. Unreacted linker was removed by buffer exchange to PBS using Amicon centrifugal filters. Azide-functionalized DNA was added at 5-fold molar excess to the DBCO-nanobody and reacted overnight at 4 °C. Nanobody-DNA conjugates were purified by anion exchange chromatograhpy using an ÄKTA Pure system with a Resource Q 1-ml column.

**Fluorophore / Antibody conjugation:** CF680 conjugated secondary F(ab')1 fragments were generated via succinimidyl-ester reaction. 100 µg anti rabbit F(ab')1 fragment (Jackson ImmunoResearch, 111-007-008) were incubated (1h RT) with a five-fold excess of succinimidyl esters of CF 680 (Biotinum, 92139) in $NaHCO_3$ (50 mM, pH 8.1). Unbound dye was removed with Nap-5 Sephadex G 25 columns (GE Healthcare). Labelled antibody was eluted from the column with PBS and stored with 50% glycerol (v/v) and 2% BSA (w/v) at -20°C.

**Cell Culture:** NIH 3T3 cells (Figure S2: only imagers P1, P3) and COS-7 cells (all other experiments) were cultured and seeded into µ-Slides (DNA-PAINT: 8 Well #1.5, Ibidi) or 24 mm coverslips (dSTORM: #1.5, Roth PK 26.1) with high glucose Dulbecco's modified Eagle's medium (DMEM) supplemented with 10% (v/v) heat-inactivated bovine calf serum (BCS, NIH 3T3) or fetal calf serum (FBS, COS-7) containing 50 U / ml penicillin and 50 µg / ml streptomycin. NIH 3T3 cells were cultured for 24 h without serum before fixation (serum starvation).

**Immunocytochemistry:** Cells were fixed with methanol (Figure S2: only imagers P1, P3) or glutaraldehyde (all other experiments). Methanol fixation: Cells were fixed with methanol (-20°C, 5 min) and then washed 3x with PBS. Glutaraldehyde fixation: Cells were pre-extracted with 0.1% glutaraldehyde and 0.25% Triton X-100 in PEM buffer (80 mM PIPES, 5mM EGTA, 2mM $MgCh_2$, pH 6.8) for ~ 30 s (37°C), followed by a fixation with 0.5% glutaraldehyde and 0.25% Triton X-100 in PEM buffer for 10 min (37°C). Cells were kept throughout the fixation on a pre-heated metal pad or in a cell culture incubator (all 37°C). Cells were then washed with PBS and incubated with 0.1% $NaBH_4$ in PBS (7 min, RT) before washing 3x with PBS. For Immunocytochemistry, cells were blocked for 1 h (RT) in blocking buffer (0.23% [v/v] Triton X-100, 385 mM NaCl in 15 mM sodium phosphate buffer pH 7.4, supplemented with 30% [v/v] goat serum) before incubation with the specific primary antibody (1 h, RT). Cells were then washed 3x with PBS and incubated with secondary nanobodies (DNA-PAINT) or F(ab')1 fragments (dSTORM) in blocking buffer (1 h, RT). Triple-color samples were stained sequentially: first with the primary and secondary for α-tubulin (ms) and clathrin (rb), washed 3 x with PBS and then with the primary and secondary for vimentin (rb). Samples were post-fixed with 4 % (w / v) PFA and 4 % (w / v) sucrose in PBS (RT) and quenched with 50 mM Ammoniumcloride in PBS for 5 minutes (RT). 150 nm gold nanoparticles (Cytodiagnostics , G-150-20) were diluted 1:10 in PBS and applied to the imaging chambers. Gold particles were allowed to settle onto the bottom (5 min RT) and washed with PBS.

**Antibodies:** The following antibodies were used: Rabbit anti clathrin heavy chain (Abcam, ab21679), mouse anti α-tubulin (Synaptic Systems, 302211), rabbit anti vimentin (Abcam, ab92547), R3-/R4-conjugated nanobodies anti rabbit and R6-conjugated nanobodies anti mouse: custom conjugations of the DNA oligo (metabion) with nanobodies (NanoTag) as decribed above. F(ab')1 Alexa 647 anti-mouse (Jackson ImmunoResearch, 115-607-003), F(ab')1 CF680 anti rabbit: custom conjugation of CF680 (Biotium, 92139) with anti-rabbit F(ab')1 fragments (Jackson ImmunoResearch, 111-007-008) as described above.

**Spectral Demixing microscope setup:** Images were acquired using the N-STORM system (version 3, Nikon) consisting of an inverted fluorescence microscope (Eclipse Ti, Nikon) controlled by NIS-Elements (Nikon, version 5.21.02), equipped with a 100x oil-immersion objective (NA: 1.49, SR HP Apochromat TIRF, Nikon) and a 1.5 magnification lens. We used the back-illuminated sCMOS camera (Prime 95B, Photometrics) using the gain settings (DNA-PAINT: 16 bit 'combined gain', dSTORM: 12 bit 'high gain' for the weaker localization intensities) and a 647 nm laser (80 mW at fiber output power, ~3 kW/cm$^2$ on the sample). Images were acquired with the following filter set: Nikon TIRF-Quad Filterset 405/488/561/647: Laser Clean-up ZET405/488/561/647 (F69-647, AHF), beam splitter zt405/488/561/647rpc flat (F73-646, AHF), emission filter ZET405/488/561/647 TIRF (F57-407, AHF). Optosplit III: emission splitter T 720 LPXR (for SD-DNA-PAINT, F43-721W, AHF), emission splitter 700-DCXXR (AHF).

**SD-DNA-PAINT Image Acquisition:** In order to minimize drift, all experiments were carried out in a live-cell incubator (Okolab) at 28°C to stabilize the temperature a few degrees above the room temperature. Imagers were diluted in imaging buffers B or C as described previously [9]. Buffer B was used for all experiments with the imagers P1 and P3 and for all origami experiments (5 mM Tris-HCl, 10 mM MgCl2 and 1 mM EDTA, pH 8.0). Buffer C was used for all cell experiments with R3S, R4S or R6S (1 × PBS, 1 mM EDTA, 500 mM NaCl, pH 7.4). Imagers were applied in an equimolar ratio to a total concentration of 1 nM. We acquired movies with 20.000 – 50,000 frames (Table 3) with the 647 nm laser at maximum power ( ~3 kW/cm$^2$ on sample) in HILO mode (highly inclined and laminated optical sheet) with engaged auto focus system (PFS, Nikon).

**SD-dSTORM image acquisition:** Images were acquired with the system described above. For SD-dSTORM samples were mounted with dSTORM imaging buffer (50 mM Tris/HCl, 10 mM NaCl, 100 mM β-mercaptoethylamine [MEA, Sigma 30070] and 0.5 mg/ml glucose oxidase

[Sigma G2133], 40 µg/ml catalase [Sigma C100] and 10% (w/v) glucose, pH 8) onto 100 µl spherical void slides (Carl Roth, H884.1) [13].

**Single-molecule localization and drift correction:** The open-source ImageJ [26] plugin ThunderSTORM1.3 [27] was used to localize blinking events from single imager/docking strand hybridizations and to perform bead-based drift correction. Single blinking events were fitted with the 'integrated Gaussian' and 'weighted least squares' method of ThunderSTORM with a fitting radius of 4 pixels (292 nm) and an initial sigma of 1.5 pixels (110 nm). Drift was calculated from the track of the gold beads in the sample (intensity threshold = 30,000) and then applied to all localizations in the image with using 'smoothingbandwith' = 0.00025. Further details are provided in the configuration files (Supplementary Information).

**Spectral Demixing:** The emission of all single-molecule events is split by the OptoSplit III into the short and long wavelength channel according to the emission spectrum of the dye attached to the imager. Each single-molecule event therefore has a localization in both channels (sides) of the camera. Depending on the emission spectra of the dye, the intensity values in each channel have a different ratio. We used our recently published open-source software tool SD-Mixer2 (available at GitHub: https://github.com/gtadeus/sdmixer2 [28]) to pair these localizations and for color-assignment of the localizations [16]. Initially, we performed drift-correction using ThunderSTORM with the entire localization files. The drift-corrected localization files were then converted into SD-Mixer file format (custom Python script available on GitHub: https://github.com/ngimber/Converter_ThunderSTORM_SDmixer [29]). The distance between the two paired localizations (offset) was initially determined manually within the raw data. The SD-Mixer2 was then used with this initial offset value to identify the localization pairs and to assign the colors to each of them. Specifically, localization pairs within each frame of the image sequence were identified using the SD-Mixer2 with the 'offset optimization' enabled and a search range of 150 nm. 'Grouping' of localizations from successive frames was enabled with a radius of 3 pixels (219 nm). Color assignment was done based on the 2D intensity histogram (Figure 3a, 4a). The binary masks for color assignment were manually drawn on top of the channel-specific 2D intensity distribution from single-color experiments, in order to maximize inclusion of localizations and to minimize color crosstalk (Figure 3b, 4b). The binary masks (Supplementary Information) were loaded into the SD-Mixer2 to filter localization pairs based on the position in the 2D intensity histogram. Finally, each localization pair was assigned to a certain color according to their position within the binary masks.

**Intensity-weighted multichannel registration and rendering:** For accurate multichannel registration and to use the single-molecule emissions from both channels (short and long wavelength, Figure 1b) we developed a novel intensity-weighted multichannel registration procedure. Registration and unwarping of the distortions was performed using a custom-written Python script that is available at GitHub: https://github.com/ngimber/SD-DNA-PAINT [30]. Specifically, the offset values (yellow vectors in Figure 1b) between short and long channel localizations were binned into a 2D matrix (1 nm bin size). We used the median from all localization within each bin as the local offset value for the correction. That way the correction was based on several localizations per bin. Therefore, errors of this correction were below the localization precision (NeNA) of the individual pairs (Figure 2e, S5). After this registration, we weighted the deviation of each localization pair from a channel-specific reference point by the intensity values in each individual channel using the following formula (equation 1):

$$PC_n = \frac{((PS_n - \frac{\sum_{i=1}^{n} PS_n}{n}) \times IS_n) + ((PL_n - \frac{\sum_{i=1}^{n} PL_n}{n}) \times IL_n)}{IS_n + IL_n} \quad (1)$$

*PC = corrected position; PS = short channel position; PL = long channel position; IS = short channel intensity; IL = long channel intensity; n = localization index*

Images were then reconstructed from the reassigned localizations with ThunderSTORM using a pixel size of 2 nm (origamis) or 5 nm (cells). The images were rendered using a Gaussian blur with sigma set to the experimentally determined NeNA precision (Figure 2e).

**Crosstalk and proportion of included localizations:** Statistics on the crosstalk and the proportion of 'included' localizations were determined based on ground truth data from single-color DNA-PAINT experiments (clathrin: R3S ATTO 700, R4S ATTO 680; α-tubulin: R6S ATTO 655). Single-color experiments were acquired in the SD-mode including grouping. We excluded regions with gold particles and used the dual- and triple-color spectral filters from Figure 3a and 4a to test how many localizations fall into the correct color channel ('included' localizations) and how many localizations fall into a wrong color channel ('crosstalk'). Crosstalk and the proportion of 'included localizations' were calculated per image and plotted as mean ± SEM over all images.

**Localization precision:** DNA origami structures with a 3 x 4 docking strand arrangement (20 nm spacing) were used to determine the localization precision of the SD microscope system. Single-color DNA-PAINT experiments were acquired in the SD-mode, including all processing steps from Fig. S2a (if not indicated differently) except of 'grouping' (as grouping is a step within the NeNA procedure). Gold beads used for drift correction were excluded and origamis were detected with the "Pick Similar" function of the Picasso software [18]. The NeNA precision per image was calculated with Picasso based on the selected docking sites and plotted as mean ± SEM from all images, N = 4 - 6 images.

**Nearest neighbor distance analysis:** In order to quantify the accuracy of the multichannel registration, DNA origami structures with a 3 x 4 docking strand arrangement (20 nm spacing) were imaged and processed as described above, including multichannel registration and intensity-weighted localization reassignment. Random background localizations were removed by Voronoi tesselation [31] and thresholding of the localization densities, both with a custom-written Python script (available on GitHub: https://github.com/ngimber/SMLM_VoronoiTesselation [32]). Gold beads were then excluded before calculating the nearest neighbor distributions between two image channels using a custom-written Python script (available on GitHub:

https://github.com/ngimber/SMLM_NearestNeighbor [33]). Randomized control images were generated by introducing a toroidal shift of 10 nm to the ‘subject channel’.

### SUPPLEMENTARY TABLES

**Supplementary Table 1.** Imager Sequences

| Name | Sequence |
|---|---|
| P1 | 5’ CTAGATGTAT 3’ - Dye |
| P3 | 5’ GTAATGAAGA 3’ - Dye |
| R3S | 5’ GAGAGAGAAA 3’ - Dye |
| R4S | 5’ TGTGTGTTT 3’ - Dye |
| R6S | 5' TTGTTGTTT 3’ - Dye |

**Supplementary Table 2.** Docking Sequences

| Name | Sequence |
|---|---|
| P1 | 5’ TTATACATCTA 3’ |
| P3 | 5’ TTATCTACATA 3’ |
| R3 | 5’ CTCTCTCTCTCTCTCTCTC 3’ |
| R4 | 5’ ACACACACACACACACACA 3’ |
| R6 | 5’ AACAACAACAACAACAACAA 3’ |

**Supplementary Table 3.** Experimental settings

| Figure | Buffer | Mode | Imager | Frames | Exposure | Processing |
|---|---|---|---|---|---|---|
| 1c | C | Single-color | R3S 700 1nM / R6S 655 1nM | 20,000 | 100 ms | L, D, P, G |
| 1d | C | Multicolor | R3S 700 0.5nM + R6S 655 0.5nM | 30,000 | 100 ms | L, D, P, G, F, M, W, R |
| 2b: 655 | C | Single-color | R3S 655 1nM | 30,000 | 100 ms | L, D, P, G, M, W, R |
| 2b: 680 | C | Single-color | R4S 680 1nM | 20,000 | 100 ms | L, D, P, G, M, W, R |
| 2b: 700 | C | Single-color | R6S 700 1nM | 30,000 | 100 ms | L, D, P, G, M, W, R |
| 2d: 655 | B | Single-color | R3S 655 1nM | 20,000 | 100 ms | L, D, P, G, M, W, R |
| 2d: 680 | B | Single-color | R4S 680 1nM | 20,000 | 100 ms | L, D, P, G, M, W, R |
| 2d: 700 | B | Single-color | R6S 700 1nM | 20,000 | 100 ms | L, D, P, G, M, W, R |
| 2e | B | Single-color | R3S 700 1nM / R4S 680 1nM / R6S 655 1nM | 20,000 | 100 ms | L, D, P, M, W |
| 3a | C | Single-color | R3S 700 1nM / R6S 655 1nM | 20,000 | 100 ms | L, D, P, G |
| 3b | C | Single-color | R3S 700 1nM / R6S 655 1nM | 20,000 | 100 ms | L, D, P, G, F |

| | | | | | | |
|---|---|---|---|---|---|---|
| 3c,d | C | Multicolor | R3S 700 0.5nM + R6S 655 0.5nM | 30,000 | 100 ms | L, D, P, G, F, M, W, R |
| 4a | C | Single-color | R3S 700 1nM / R4S 680 1nM / R6S 655 1nM | 20,000 | 100 ms | L, D, P, G |
| 4b | C | Single-color | R3S 700 1nM / R4S 680 1nM / R6S 655 1nM | 20,000 | 100 ms | L, D, P, G, F |
| 4c | C | Multicolor | R3S 700 0.3nM + R4S 680 0.3nM + R6S 655 0.3nM | 30,000 | 100 ms | L, D, P, G, F, M, W, R |
| 4d,e | B | Multicolor | R5S 700 0.3nM + R5S 680 0.3nM + R6S 655 0.3nM | 50,000 | 100 ms | L, D, P, G, F, M, W, R |
| S1c-f | B | Single-color | R3S 700 1nM / R4S 680 1nM / R6S 655 1nM | 20,000 | 100 ms | L, D, P, G, M, W, R |
| S2b | C | Single-color | R3S 700 1nM / R4S 680 1nM / R6S 655 1nM | 20,000 | 100 ms | as indicated |
| S2c: 655 | B | Single-color | R3S 655 1nM | 30,000 | 100 ms | as indicated |
| S2c: 680 | B | Single-color | R4S 680 1nM | 20,000 | 100 ms | as indicated |
| S2c: 700 | B | Single-color | R6S 700 1nM | 30,000 | 100 ms | as indicated |
| S2d | B | Single-color | R3S 700 1nM / R4S 680 1nM / R6S 655 1nM | 20,000 | 100 ms | L, D, P, M, (W if indicated) |
| S3a | C | Single-color | R3S 700 1nM / R6S 655 1nM | 20,000 | 100 ms | L, D, P, G, F |
| S3b | C | Single-color | R3S 700 1nM / R4S 680 1nM / R6S 655 1nM | 20,000 | 100 ms | L, D, P, G, F |
| S3c | B | Multicolor | R3S 700 1nM / R4S 680 1nM / R6S 655 1nM | 50,000 | 100 ms | L, D, P, G, F, M, W, R |
| S4a,b | C | Multicolor | - | 30,000 | 30 ms | L, D, P, G, F, M, W, R |
| S4c | C | Multicolor | - | 30,000 | 30 ms | L, D, P, F, M, (W if indicated) |
| S5a | B | Multicolor | R6S 700 1nM / R6S 680 1nM / R6S 655 1nM | 50,000 | 100 ms | L, D, P, G, F, M, W |
| S5b | B | Multicolor | R3S 700 1nM / R4S 680 1nM / R6S 655 1nM | 50,000 | 100 ms | L, D, P, G, F, M, W |

L = localization, D = drift correction, P = pairing, G = grouping, F = color filtering, M = multichannel registration, W = intensity-weighting, R = rendering.
