## Supplementary figures and images for "Simultaneous multicolor DNA-PAINT without sequential fluid exchange using spectral demixing"

### ch1_20200827_680.png

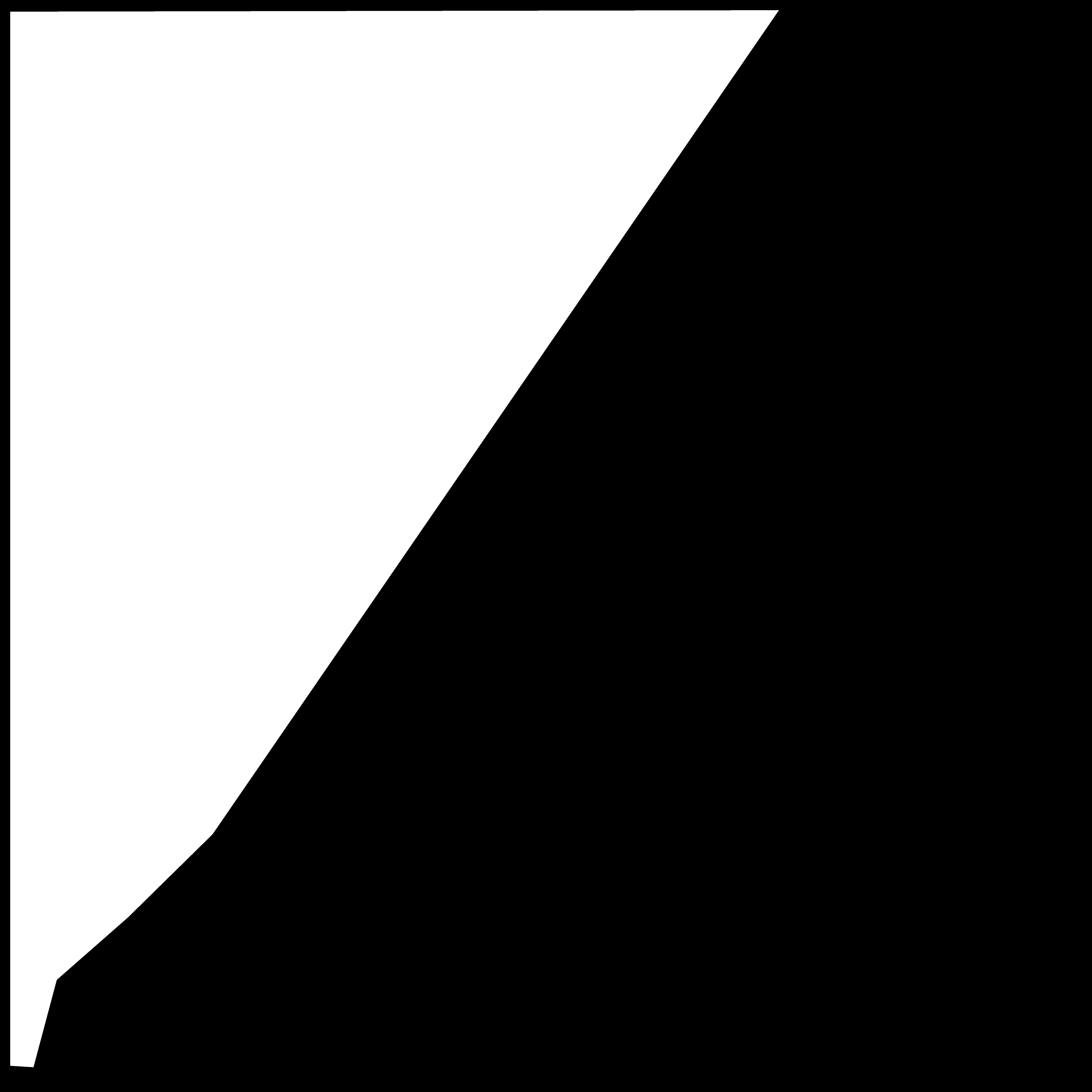

### ch1_mask700_dual_final.png

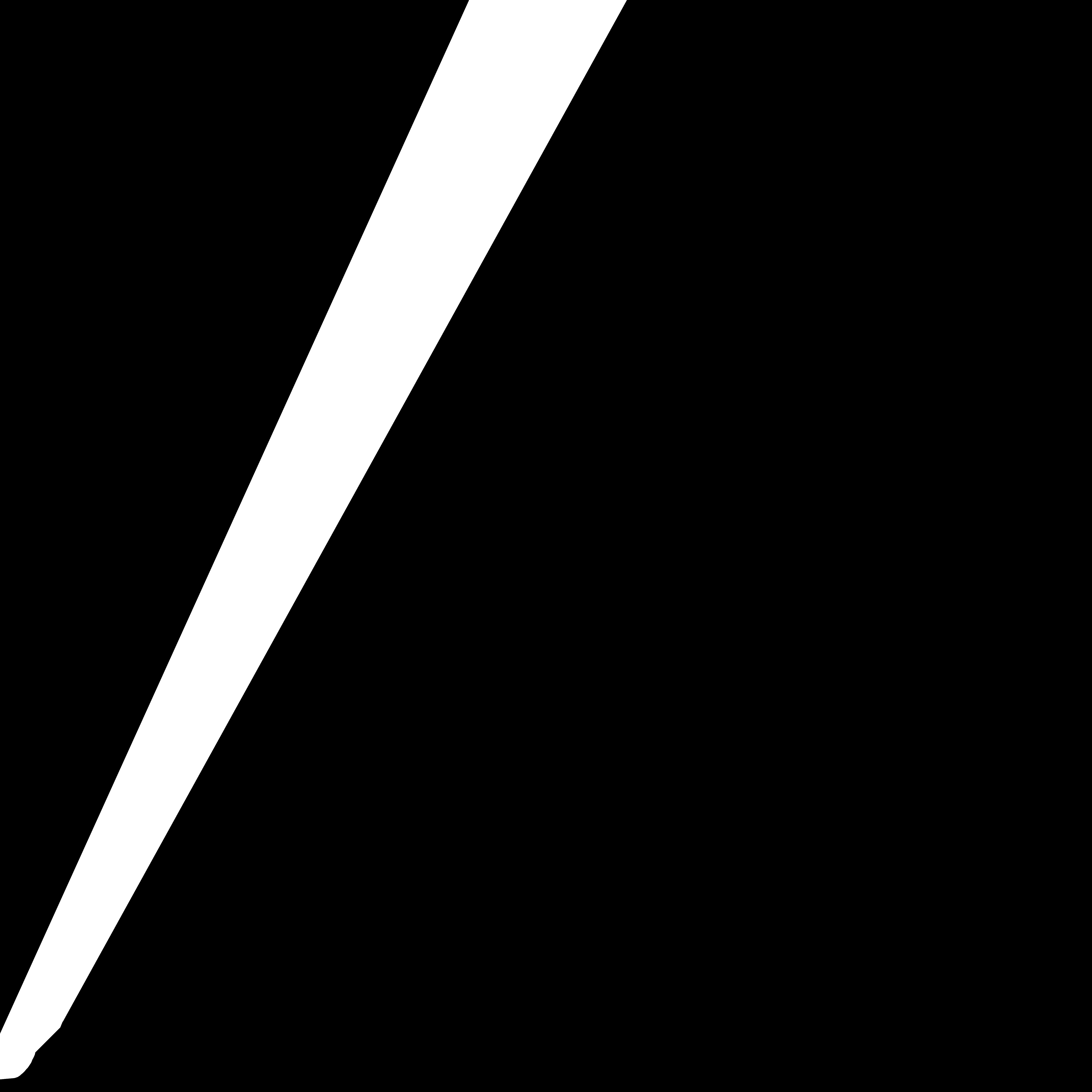

### ch2_20200827_647.png

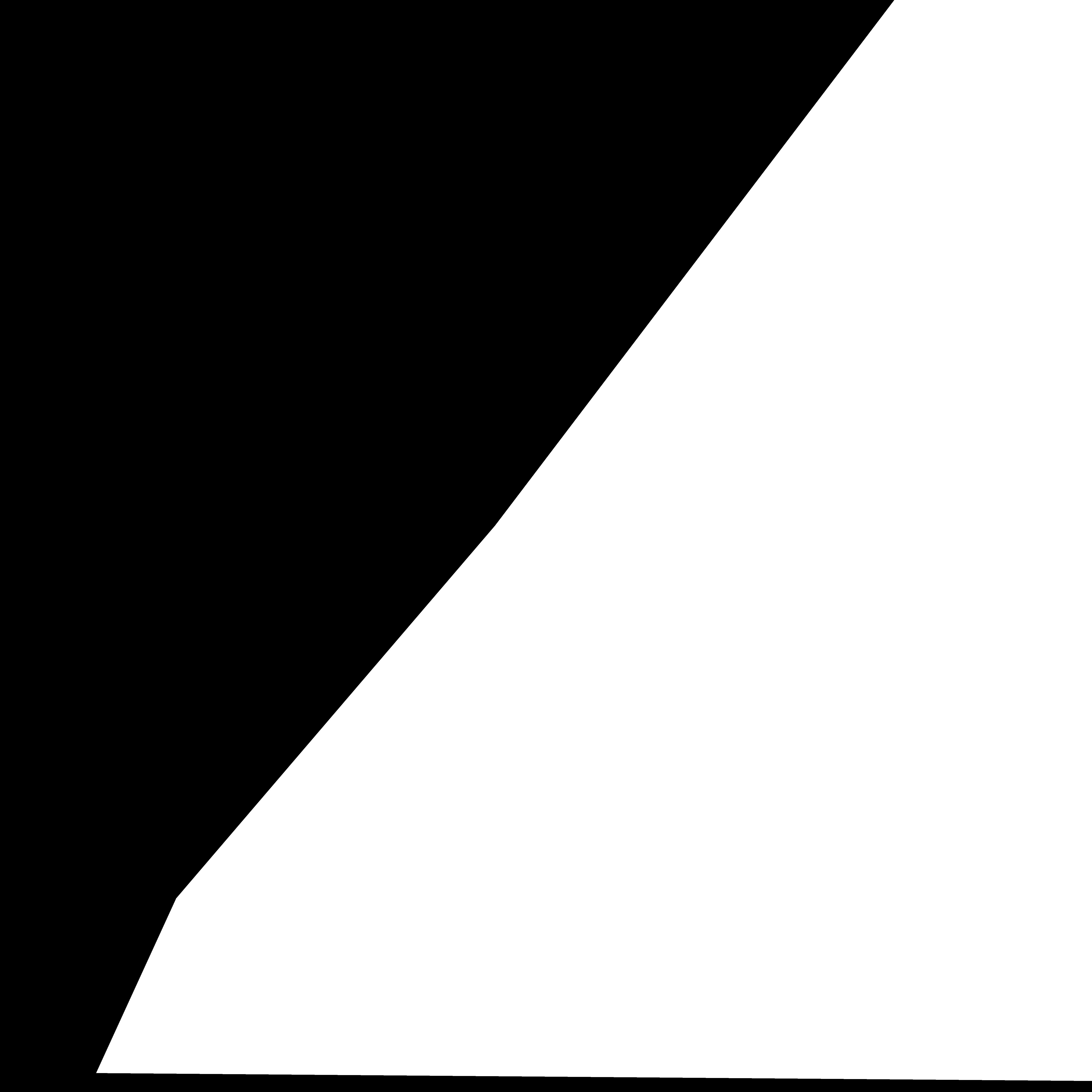

### ch2_mask655_dual_final.png

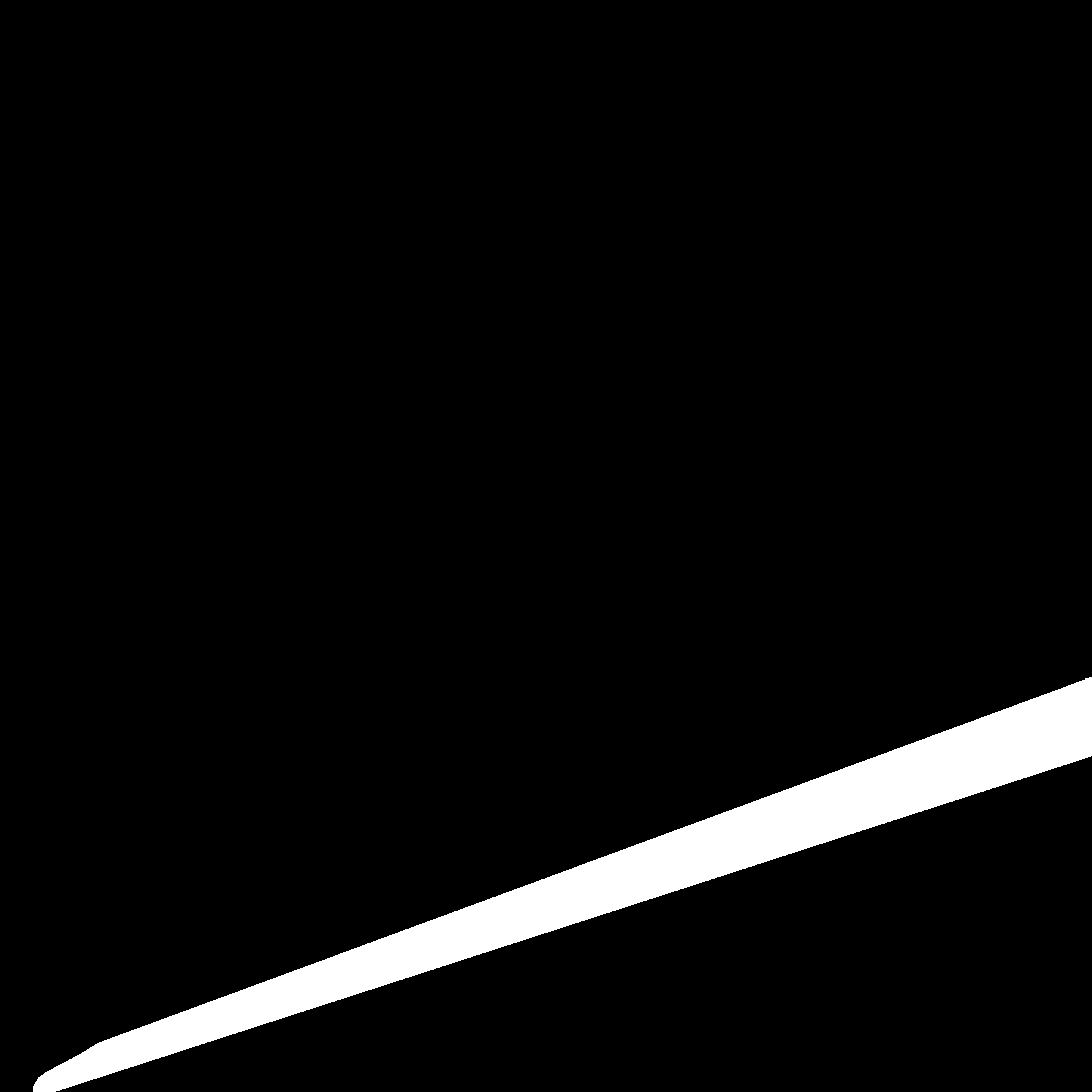

### ch3_mask700_tripple_final.png

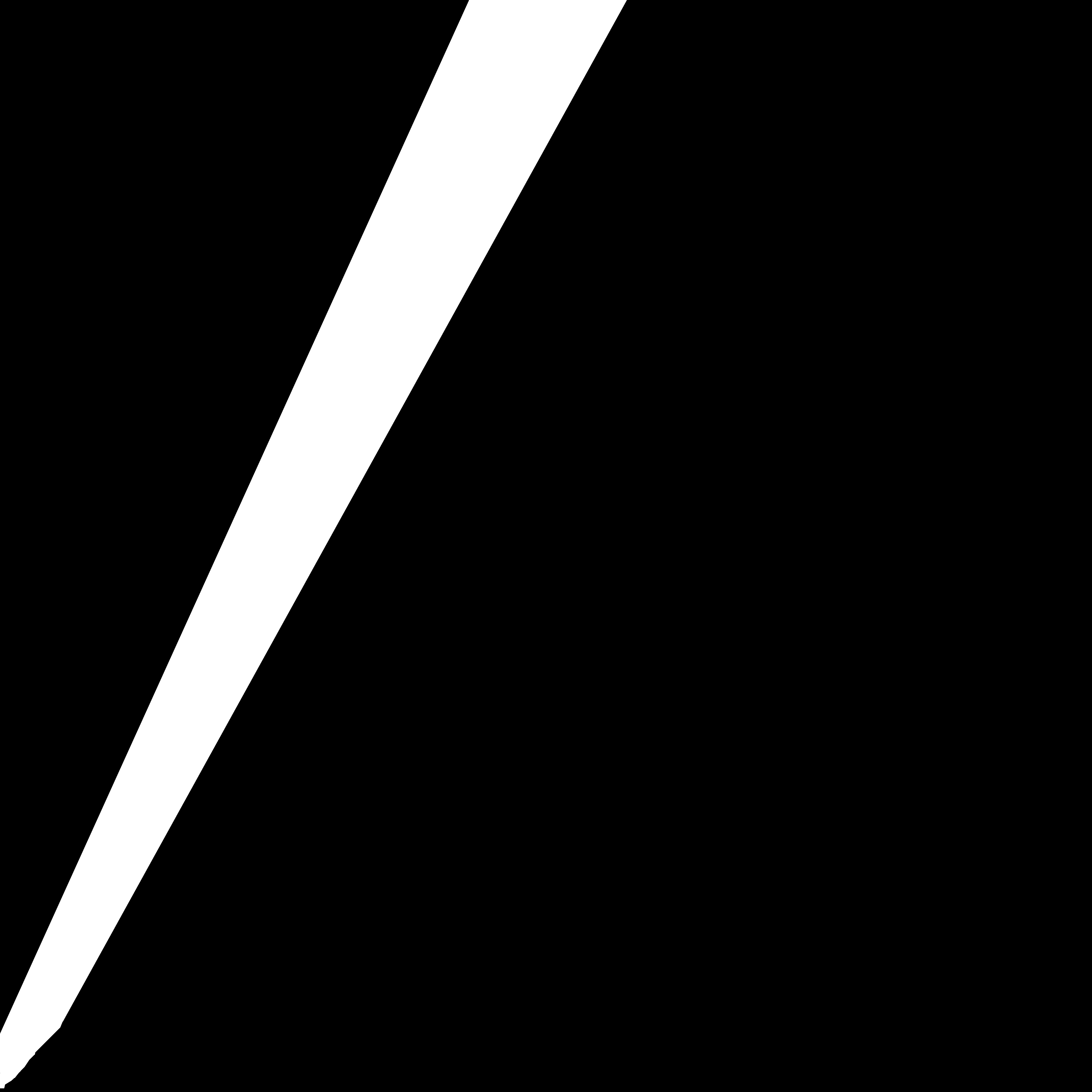

### ch4_mask680_tripple_final.png

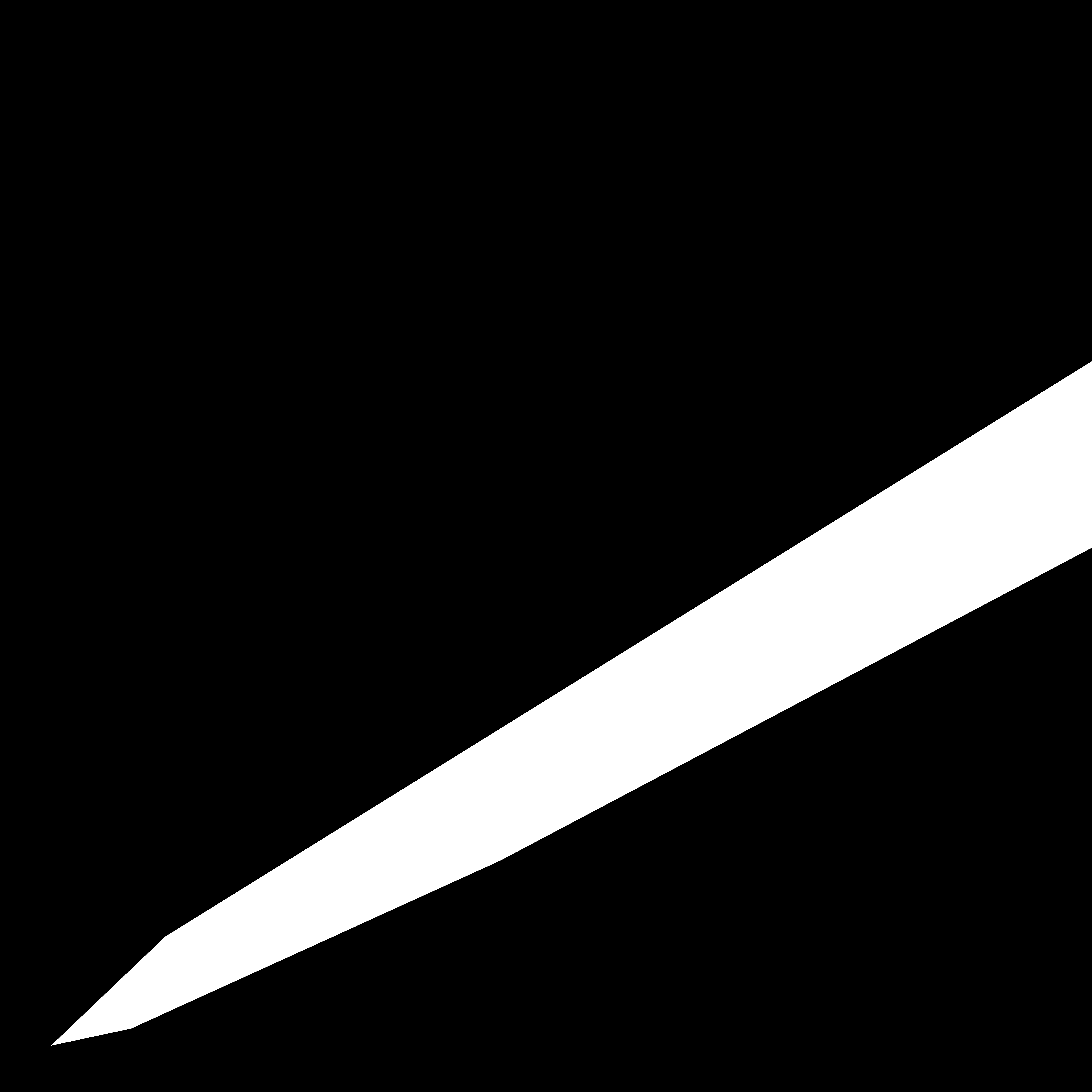

### ch5_mask655_tripple_final.png

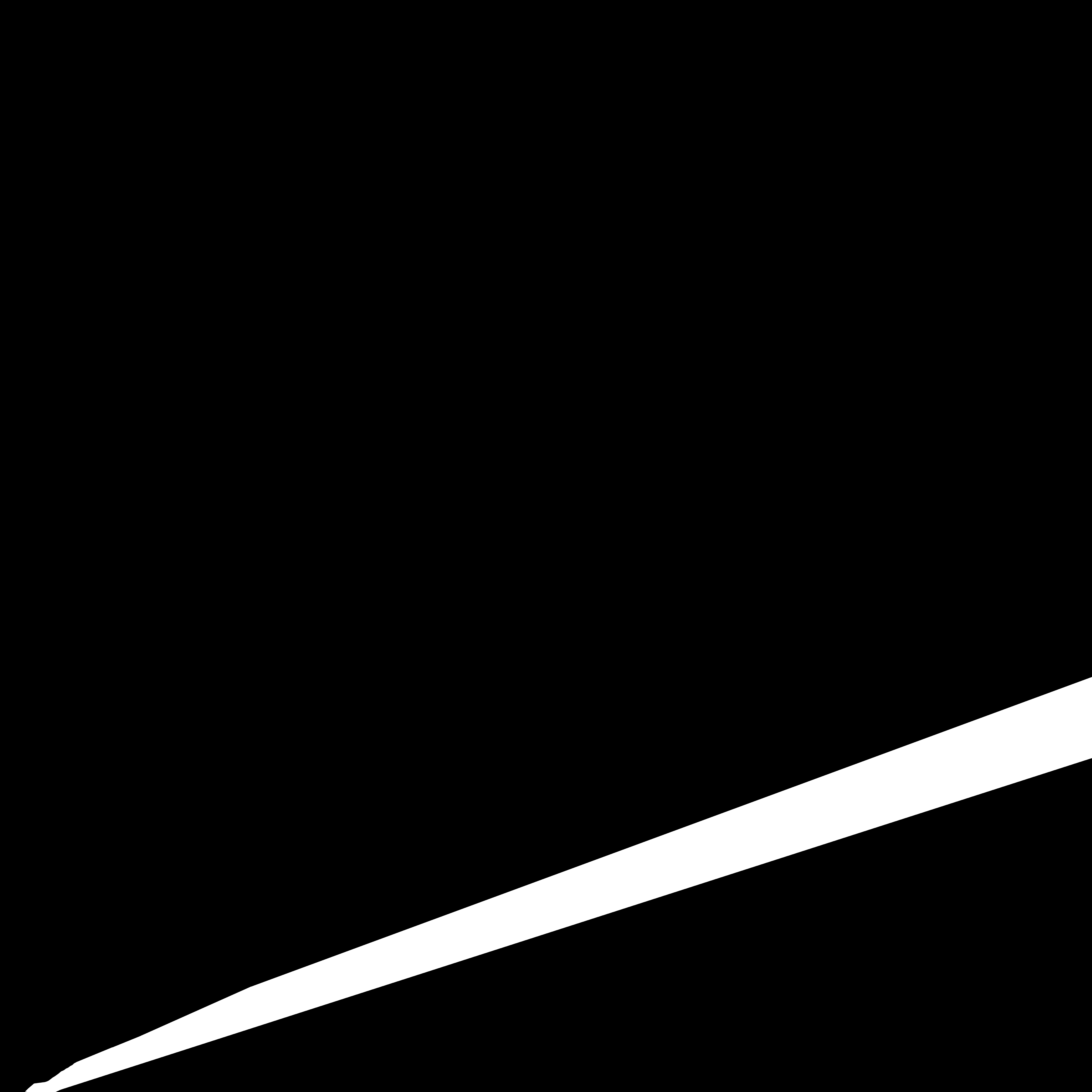
